# PI 3-kinase delta enhances axonal PIP_3_ to support axon regeneration in the adult CNS

**DOI:** 10.1101/787994

**Authors:** Amanda C. Barber, Rachel S. Evans, Bart Nieuwenhuis, Craig S. Pearson, Joachim Fuchs, Amy R. MacQueen, Susan van Erp, Barbara Haenzi, Lianne A. Hulshof, Andrew Osborne, Raquel Conceicao, Tasneem Z. Khatib, Sarita S. Deshpande, Joshua Cave, Charles ffrench-Constant, Patrice D. Smith, Klaus Okkenhaug, Britta J. Eickholt, Keith R. Martin, James W. Fawcett, Richard Eva

**Author notes:** These authors contributed equally (Amanda C. Barber, Rachel S. Evans, Bart Nieuwenhuis).

## Abstract

Peripheral nervous system (PNS) neurons support axon regeneration into adulthood, whereas central nervous system (CNS) neurons lose regenerative ability after development. To better understand this decline whilst aiming to improve regeneration, we focused on phosphoinositide 3-kinase (PI3K) and its product phosphatidylinositol(3,4,5)-trisphosphate (PIP3). We demonstrate that adult PNS neurons utilise two catalytic subunits of PI3K for axon regeneration: p110α and p110δ. However in the CNS, axonal PIP3 decreases with development at the time when axon transport declines and regenerative competence is lost. Overexpressing p110α in CNS neurons had no effect, however expression of p110δ restored axonal PIP3 and increased regenerative axon transport. p110δ expression enhanced CNS regeneration in both rat and human neurons and in transgenic mice, functioning in the same way as the hyperactivating H1047R mutation of p110α. Furthermore, viral delivery of p110δ promoted robust regeneration after optic nerve injury, demonstrating potential for translational development. These findings establish a deficit of axonal PIP3 as a key reason for intrinsic regeneration failure and demonstrate that native p110δ facilitates axon regeneration by functioning in a hyperactive fashion.

## Introduction

Adult central nervous system (CNS) neurons have a weak capacity for axon regeneration, meaning that injuries in the brain, spinal cord and optic nerve have devastating consequences (Curcio & Bradke, 2018; He & Jin, 2016). In most CNS neurons, regenerative capacity is lost as axons mature, both *in vitro* (Goldberg et al, 2002; Koseki et al, 2017) and *in vivo* (Kalil & Reh, 1979; Wu et al, 2007). Conversely, peripheral nervous system (PNS) neurons maintain regenerative potential through adult life. This is partly because PNS neurons mount an injury response in the cell body (Puttagunta et al, 2014; Smith & Skene, 1997; Ylera et al, 2009), and also because PNS axons support efficient transport of growth promoting receptors, whilst many of these are selectively excluded from mature CNS axons (Andrews et al, 2016; Franssen et al, 2015; Hollis et al, 2009a; Hollis et al, 2009b). Studies into intrinsic regenerative capacity have implicated signalling molecules, genetic factors and axon transport pathways as critical regeneration determinants (Blackmore et al, 2012; Eva et al, 2017; Fagoe et al, 2015; Hervera et al, 2019; Park et al, 2008; Weng et al, 2018). This leads to a model where axon growth capacity is controlled by genetic and signalling events in the cell body, and by the selective transport of growth machinery into the axon to re-establish a growth cone after injury.

In order to better understand the mechanisms regulating axon regeneration from within the axon and the cell body, we focused on the class I phosphoinositide 3-kinases (PI3Ks). These enzymes mediate signalling through integrins, growth factor and cytokine receptors by producing the membrane phospholipid PIP3 from PIP2 (phosphatidylinositol(3,4,5)-trisphosphate from phosphatidylinositol(4,5)-bisphosphate). Class 1 PI3Ks comprise 4 catalytic isoforms called p110α, β, γ and δ, with distinct roles for some of these emerging in specific cell populations (Bilanges et al, 2019). The p110α and p110β isoforms are ubiquitously expressed, whilst p110δ and p110γ are highly enriched in leukocytes (Hawkins & Stephens, 2015). In neurons, p110δ has been shown to be required during regeneration of dorsal root ganglion (DRG) neurons in the PNS (Eickholt et al, 2007) and p110α mediates axon growth during chick development (Hu et al, 2013) whilst p110β and p110γ have not been studied.

The class I phospholipids are strongly implicated in the regulation of regenerative ability because transgenic deletion of PTEN, an enzyme which opposes PI3K by converting PIP3 back to PIP2, promotes CNS regeneration (Geoffroy et al, 2015; Liu et al, 2010; Park et al, 2008), whilst inhibition of a negative feedback to this pathway similarly enhances regrowth (Al-Ali et al, 2017). These findings indicate a pro-regenerative role for PIP3, although this molecule has not been directly studied in adult CNS axons. Highly localised axonal PI3K activity contributes to axonal polarity in developing hippocampal axons (Shi et al, 2003), whilst spatially segregated PI3K activity is required at the growth cone of PNS axons to elicit rapid axon growth (Zhou et al, 2004). We reasoned that CNS regenerative failure might be associated with a developmental decline in axonal PIP3, and wondered whether specific PI3K isoforms were required to yield sufficient axonal PIP3 to enable regeneration.

We investigated the class I PI3K isoforms and found that both p110α and p110δ are required for PNS axon regeneration, and that p110δ is specifically required within the axon. In CNS neurons we found PIP3 was sharply downregulated with development, diminishing in the axon at the time when axon transport and regeneration also decline. We attempted to restore PIP3 through overexpression of either p110α or p110δ, however only p110δ led to elevated axonal PIP3. Importantly, by introducing the hyperactivating H1047R mutation into p110α we found that it could mimic the effect of p110δ, with expression of either facilitating axon regeneration. This suggests that regeneration is hindered by low activation of PI3K in mature CNS axons.

Furthermore, transgenic expression of p110δ or p110α^H1047R^ in adult retinal ganglion cell (RGC) neurons led to enhanced survival and axon regeneration after optic nerve crush, whilst viral expression of p110δ led to stronger regeneration. Importantly, overexpression of p110δ has both somatic and axonal effects, enhancing the axonal transport of growth machinery and signalling through ribosomal S6 in the cell body. These findings demonstrate a deficit of axonal PIP3 as a novel reason for intrinsic regenerative failure, whilst establishing that native p110δ functions in a hyperactive fashion to enable CNS axon regeneration. Our results emphasize the importance of elevating growth-promoting pathways in the axon as well as the cell body, in order to stimulate axon regeneration.

## Results

### Gene expression of p110 isoforms in the nervous system

Because localised axonal activation of PI3K is essential for rapid PNS axon growth (Zhou et al, 2004), we reasoned that there may be specific PI3K isoforms that support regeneration within PNS axons, and that these might be under-represented in CNS neurons. p110δ is plays a role in PNS axon regeneration (Eickholt et al, 2007), however the contribution of other isoforms has not been examined. We investigated the RNA expression of the individual p110 subunits (p110α, β, γ and δ) in four published neuronal RNAseq datasets (Figure S1), examining expression in DRG neurons during axon growth and regeneration (Tedeschi et al, 2016), in developing cortical neurons (Koseki et al, 2017) and in the Brain-RNAseq databases (http://www.brainrnaseq.org/) of individual cell types in mouse and human brain (Zhang et al, 2014; Zhang et al, 2016). In DRG neurons p110α, β and δ are expressed at all stages *in vitro*, whilst p110γ is expressed at very low levels. p110α is present at the highest levels, whilst p110δ is upregulated in development, and further upregulated upon peripheral nerve lesion (Figure S1A - C). In developing cortical neurons, p110α is again expressed at the highest levels, p110β and δ are present but not abundant, and p110γ is at very low levels. Remarkably, p110α, β and δ are all downregulated as cortical neurons mature (Figure S1D and E). The Brain-RNAseq database indicates p110α and β are the principal neuronal PI3K isoforms in both the mouse and human adult brain, with p110α again expressed at the highest levels, whilst p110δ and γ are enriched in microglia and macrophages (Figure S1F - M). These datasets indicate that p110α, β and δ are expressed in regenerative PNS neurons, but are almost absent or downregulated with maturation in CNS neurons. We therefore chose to investigate the contribution of the p110α, β and δ isoforms of PI3K to axon regeneration of DRG neurons.

### p110α and δ are required for DRG axon regeneration

We used laser axotomy to sever the axons of adult DRG neurons and measured the effect of specific inhibitors of p110α, β and δ on growth cone regeneration (Figure 1). Inhibiting either p110α or δ reduced the percentage axons regenerating, as did pan-PI3K inhibition (α, β and δ) or targeting p110α and δ together. Inhibiting p110β had no effect, but inhibiting all of the isoforms increased the time taken to develop a new growth cone (Figure 1A and B). Some inhibitors also caused uncut axons to stop growing, so we measured the extension rate of uncut axons. Treatment with p110α inhibitors, pan-PI3K or dual α / δ inhibitors led to a dramatic reduction in the percentage of uncut axons extending in the two-hour period, whilst they continued to extend in the presence of specific p110δ inhibitors (Figure 1C and D). We next examined the effect of PI3K inhibition using microfluidic compartmentalised chambers, in which axons extend through microchannels into a separate compartment from the cell bodies (Figure 1E). Inhibition of p110α or p110δ showed different effects depending on localization. Inhibition of p110δ in the axonal compartment reduced the percentage of regenerating axons, but had no effect in the somatic chamber. In contrast, the p110α inhibitor A66 reduced regeneration when applied either to the axonal or somatic compartment, and also increased the time taken to generate a new growth cone (Figure 1F). These data show that DRG neurons require p110α and δ for efficient regeneration. Axon growth and regeneration requires p110α activity in both the cell body and axon, whilst axon regeneration further relies on p110δ activity specifically within the axon.

**Figure 1.**
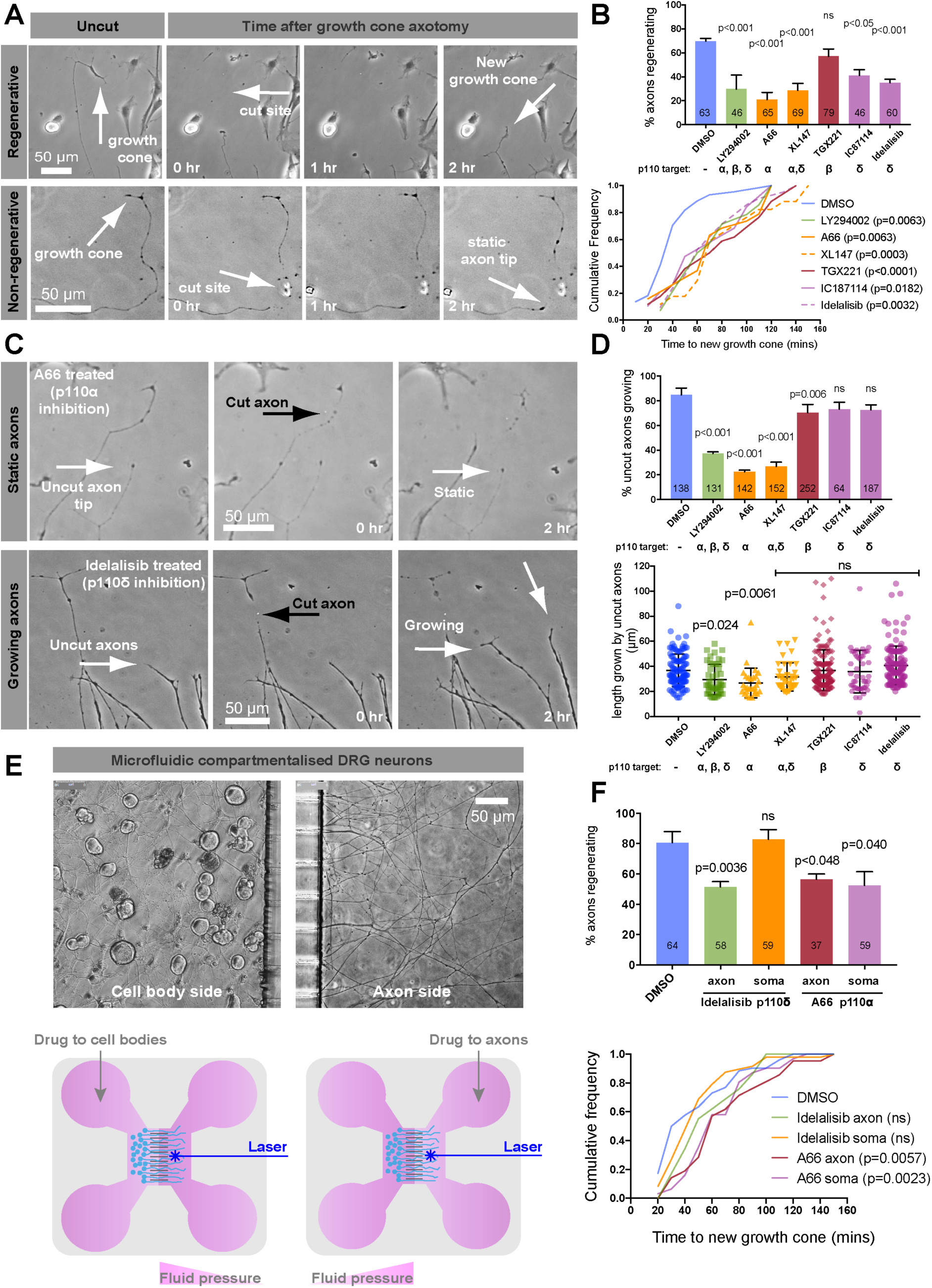
p110α and δ are required for DRG axon regeneration; p110δ functions in the axonal compartment. See also supplemental figure S1. (A) Laser injured DRG axons showing growth cone regeneration (upper panels) and regenerative failure (lower panels). Arrows as indicated. (B) Percentage of regenerating axons of DIV 1-2 adult DRG neurons in the presence of p110 inhibitors, 2 h after injury. Error bars are s.e.m. P values indicate statistical significance analysed by Fishers exact (upper graph) or Kruskal-Wallis test (lower graph). (C) Examples of static vs. growing uninjured axons treated with p110α or δ inhibitors as indicated. (D) Percentage of uninjured, growing axons in the presence of p110 inhibitors. Lower graph shows length grown in 2 h. Error bars are s.e.m. P values indicate significance measured by Fishers exact test (upper graph) or ANOVA with Tukey’s post-hoc analysis (lower graph). (E) Adult DRG neurons in microfluidic compartmental chambers. Cell bodies on the left side, axons extending through microchannels on the right. Lower panels are schematics. (F) Percentage of regenerating axons in microfluidic chambers. p110α or δ inhibitors were applied to either soma or axons. Error bars are s.e.m. P values indicate statistical significance as measured by Fishers exact test (upper graph) or by Kruskal-Wallis test (lower graph).

### PIP3 is developmentally downregulated in cortical neurons, but present in adult DRG axons

The RNAseq data described above (Figure S1) suggest that PI3K expression increases with maturation in DRG neurons whilst it is downregulated as cortical neurons mature. CNS regenerative failure might therefore be due to insufficient PI3K activity within the axon, and insufficient PIP3. PIP3 is implicated in axon growth but its developmental distribution has not been examined. PIP3 has previously been localised using fluorescently tagged pleckstrin homology (PH)-domain reporters, such as AKT-PH-GFP, however the principle readout of these reporters is translocation to the surface membrane, and they do not report on abundance or “steady state” distribution. In order to accurately measure PIP3 in neurons developing *in vitro* we optimised a fixation technique (Hammond et al, 2009) for antibody-based PIP3 detection on immobilised membrane phospholipids utilising an antibody widely used for biochemical assays. To validate this in neurons, we isolated DRG neurons from transgenic mice expressing AKT-PH-GFP at low levels (Nishio et al, 2007) to avoid inhibition of downstream signalling (Varnai et al, 2005). Live imaging of adult DRG neurons from these mice revealed dynamic hotpots of PIP3 at the axon growth cone (Figure S2A and Movie 1), and membrane labelling confirmed these are not regions of membrane enrichment (Figure S2B and Movie 2). Phospholipid fixation and labelling with anti-PIP3 revealed colocalization between AKT-PH-GFP and anti-PIP3 at regions within DRG growth cones (Figure S2C), as well as at hotspots and signalling platforms in non-neuronal cells from DRG cultures (Figure S2D). In addition to validating this technique, our data also confirm the presence of dynamic PIP3 in the growth cone and axons of regenerative DRG neurons. To further confirm the specificity of the stain for PIP3 we stimulated N1E cells with insulin and labelled for PIP3 in the presence or absence of the pan-PI3K inhibitor GDC-0941, detecting increased PIP3 staining after insulin stimulation alone, and not in the presence of the PI3K inhibitor (Figure S2 E, F and G).

We then labelled E18 cortical neurons at 3, 8 and 16 days *in vitro* (DIV) to detect endogenous PIP3. In immature neurons (at DIV 3) we detected high levels of PIP3 in the cell body, and particularly in the distal axon and growth cone (Figure 2A, D and E). At DIV 8 these neurons have a single rapidly growing axon; at this time there was a sharp decline in PIP3 at the cell body (Figure 2B and D), but only a small decrease in growth cone PIP3 (Figure 2B and E), indicating that during the period of rapid axon extension, neurons possess high levels of PIP3 in axonal growth cones. By DIV 16 cortical neurons have long axons which are establishing synapses, with some branches still growing. These mature neurons exhibit a sharp decline in regenerative ability and selective axon transport (Eva et al, 2017; Koseki et al, 2017). At this stage, we detected markedly reduced levels of PIP3 at the growth cone while they remained low at the cell body (Figure 2C, D and E). This indicates a global reduction in PIP3 as cortical neurons develop and demonstrates that the loss of regenerative ability coincides with a deficiency in axonal PIP3 production.

**Figure 2.**
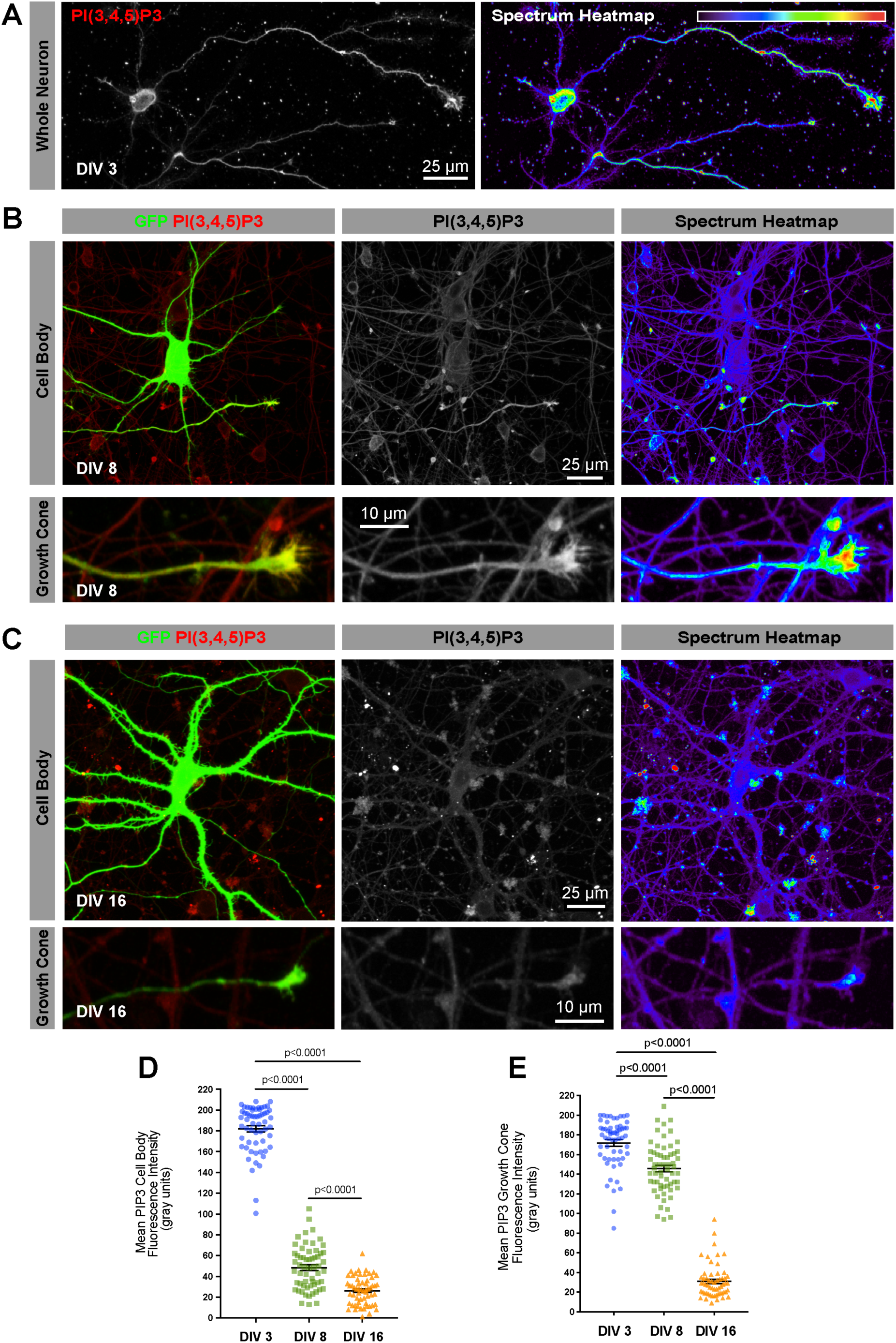
PIP3 is developmentally downregulated in cortical neurons maturing *in vitro*, first in the soma, then at the growth cone. See also supplemental figure S2 and Movies 1 and 2. (A) DIV 3 cortical neurons immunolabelled for PIP3. (B) DIV 8 cortical neuron transfected with GFP and immunolabelled for PIP3. (C) DIV 16 cortical neuron transfected GFP and immunolabelled for PIP3. (D) Somatic PIP3 quantification in the soma at increasing days *in vitro*. Error bars are s.e.m. P values are significance measured by ANOVA with Tukey’s post-hoc analysis. (E) Growth cone PIP3 quantification at increasing days *in vitro*. Error bars are s.e.m. P values indicate significance measured by ANOVA with Tukey’s post-hoc analysis.

### Expression of p110δ or p110α^H1047R^ elevates PIP3 in the soma and axon

The results above show that PI3K and PIP3 are developmentally downregulated in CNS axons as they lose their regenerative ability. In DRG neurons, which continue to express PI3K and produce PIP3, the p110α and δ isoforms are necessary for efficient axon regeneration (Figure 1). We therefore asked whether we could increase PIP3 levels and restore regeneration to CNS neurons by ectopic expression of p110α, its hyperactive H1047R variant p110α^H1047R^ (Mandelker et al, 2009) or p110δ. Previous work has shown that p110δ and p110α^H1047R^ can sustain downstream AKT activation upon overexpression in fibroblasts, acting independently of active Ras, whilst native p110α does not (Kang et al, 2006). These PI3Ks were expressed in cortical neurons at DIV 16, and PIP3 was measured by quantitative immunofluorescence. We used dual-promoter constructs expressing untagged p110 and GFP, in order to avoid potential interference of PI3K function by a protein tag.

Compared with cells expressing GFP alone, expression of p110α had no effect on PIP3 levels in DIV 16 neurons, either in the cell body or within the axons (Figure 3 A, B,C and D). In contrast, overexpression of either p110α^H1047R^ or p110δ led to a small but significant increase in PIP3 in the cell body (Figure 3, A and C), and a striking increase in PIP3 at axon growth cones (Figure 3, B and D). These findings indicate that p110δ and p110α^H1047R^ function in a hyperactive fashion to generate PIP3 in cortical neurons, with the strongest effect in growth cones. However, even overexpressed p110α does not generate PIP3, suggesting a low level of activation in mature neurons.

**Figure 3.**
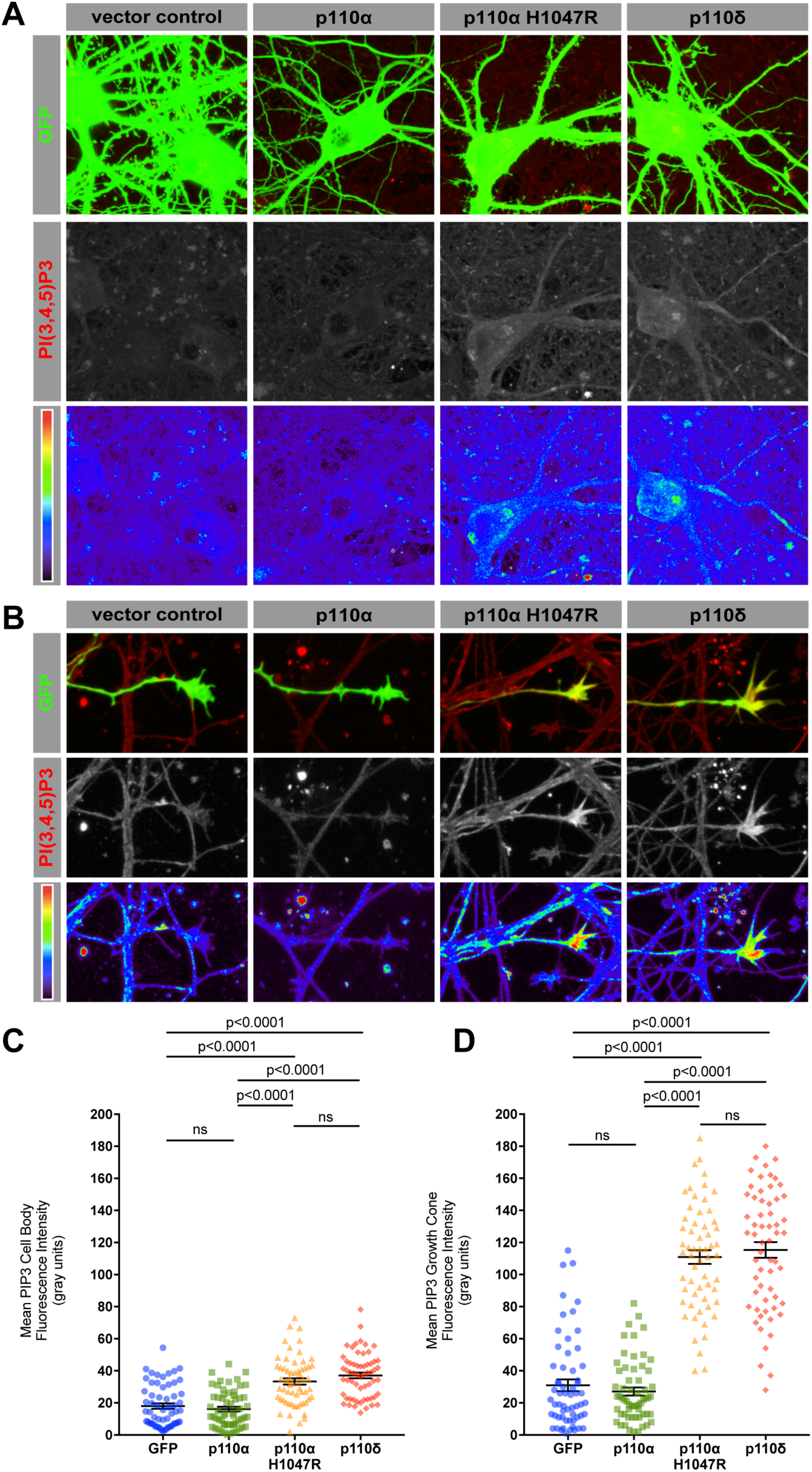
p110δ or p110α^H1047R^ expression elevates PIP3 in the soma and axon, whilst native p110α does not. (A) PIP3 immunofluorescence in the soma of DIV 16 cortical neurons expressing p110 isoforms and GFP. Spectrum heatmap shows fluorescence intensity. (B) PIP3 immunofluorescence at the axonal growth cone of DIV 16 neurons expressing p110 isoforms and GFP. Spectrum heatmap shows fluorescence intensity. (C) Quantification of PIP3 immunofluorescence in the soma. Error bars are s.e.m. P values indicate significance as measured by ANOVA with Tukey’s post-hoc analysis. (D) Graph showing PIP3 quantification of immunofluorescence at the axon growth cone. Error bars are s.e.m. P values indicate significance measured by ANOVA with Tukey’s post-hoc analysis.

### Expression of p110δ or p110α^H1047R^ increases axon and dendrite growth

We next investigated the effect of p110δ, p110α or p110α^H1047R^ overexpression on the regulation of axon growth in cortical neurons developing *in vitro*. Expression of either p110δ or p110α^H1047R^ in immature neurons at DIV 2 led to a moderate increase in axon length by DIV 4, compared with neurons expressing either GFP alone or native p110α (Figure 4A and B). p110δ or p110α^H1047R^ expression also led to a small increase in the dendrite length (Figure 4A and C). This led to an increase in axon/dendrite length ratio, with axons approximately 31.5 times longer than dendrites, while control neurons have axons 21.8 times longer than dendrites. None of the PI3K isoforms therefore affected neuronal polarisation (Figure 4A and D). We also examined the effects of PI3K overexpression on dendrite length and branching at a later developmental stage (transfecting at DIV 10, and analysing at DIV 14). Again, p110δ and p110α^H1047R^ behaved similarly, expression of either construct leading to increases in both the number of dendrite branches and the total dendrite length, compared with control-transfected neurons. In contrast, expression of native p110α had no effect (Figure 4E, F and G). The PI3K pathway is a well known regulator of cell size, so we also measured hypertrophy. Overexpression of either p110δ or p110α^H1047R^ led to an increase in cell body size, compared with either GFP or p110α (Figure 4H). To confirm downstream signalling through the PI3K pathway, we employed phosphorylation-specific immunolabelling of ribosomal S6 protein, a transcriptional regulator routinely used as a reporter of somatic signalling through the PI3K/AKT/mTOR pathway. p110δ or p110α^H1047R^ expression led to a strong phospho-S6 signal compared with GFP expressing controls, whilst expression of p110α again had no effect (Figure 4E and I). These findings confirm that hyperactive p110α^H1047R^ behaves like p110δ to trigger downstream signalling through the PI3K pathway in neuronal soma, with effects on size, dendrite branching and axon length. The results demonstrate that p110δ and p110α^H1047R^ enhance both axonal and dendritic growth whilst native p110α does not.

**Figure 4.**
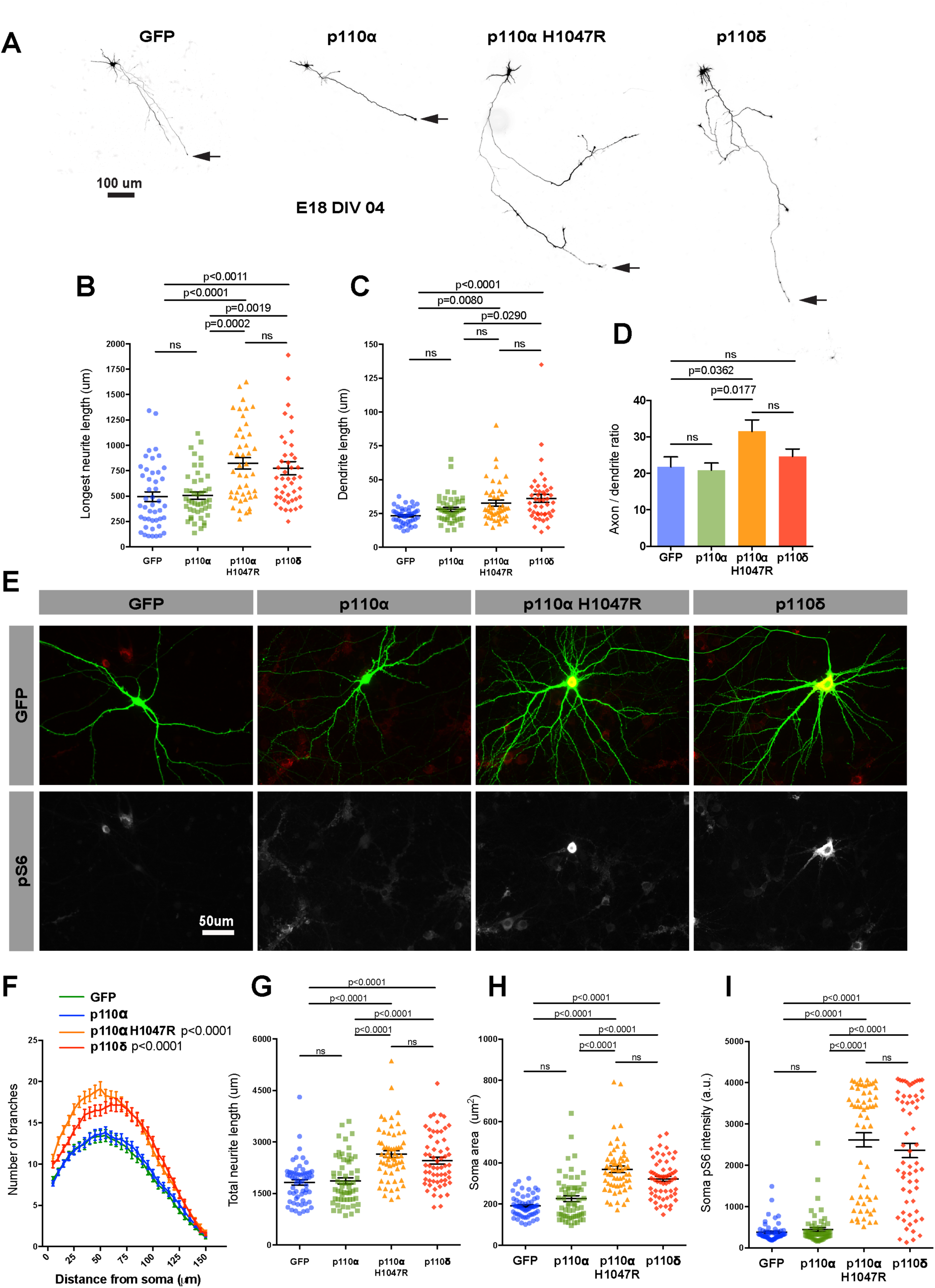
Expression of p110δ or p110α^H1047R^ increases axon and dendrite growth of cortical neurons developing *in vitro*, whilst native p110α does not. (A) DIV 4 cortical neurons expressing p110 isoforms and GFP. Arrow marks the axon tip. (B) Quantification of axon length. n = 45 neurons per group. (C) Quantification of dendrite length. n = 45 neurons per group. (D) Quantification of the axon:dendrite length ratio. n = 45 neurons per group. (E) DIV 14 cortical neurons expressing p110 isoforms and GFP, immunolabelled for pS6. (F) Sholl analysis of branches. n = 60 neurons per group. Error bars are s.e.m. P values indicate significance measured by repeated measure ANOVA with Bonferonni’s post-hoc test. (G) Quantification of the total neurite length. n = 60 neurons per group. (H) Quantification of soma area. n = 60 neurons per group. (I) Quantification of the pS6 immunofluorescence. n = 60 neurons per group. B-D and H-I, Error bars are s.e.m. P values indicate significance measured by ANOVA with Tukey’s post-hoc analysis.

### p110δ and p110α^H1047R^ promote axon regeneration of CNS neurons *in vitro*

To determine whether activation of p110 can facilitate CNS regeneration we uswed an *in vitro* model of regeneration in mature cortical neurons, comparing overexpression of either p110δ, p110α or p110α^H1047R^. In this model, axon regeneration ability is progressively lost with maturity, and molecules that promote regeneration may differ from those that promote developmental outgrowth (Koseki et al, 2017). *In vitro* laser axotomy was used to sever the axons of E18 cortical neurons cultured to DIV 15-18, at which stage they have a limited capacity for regeneration (Eva et al, 2017; Koseki et al, 2017). Expression of either p110δ or p110α^H1047R^ led to a sharp increase in the percentage of axons regenerating after axotomy compared with GFP expressing controls. p110α expression again had no effect (Figure 5A and B and Movies 3 and 4). Expression of p110δ or p110α^H1047R^ also led to an increase in the length of regenerated axons, and a trend towards shorter time of onset to regeneration compared to controls (Figure 5 C and D).

**Figure 5.**
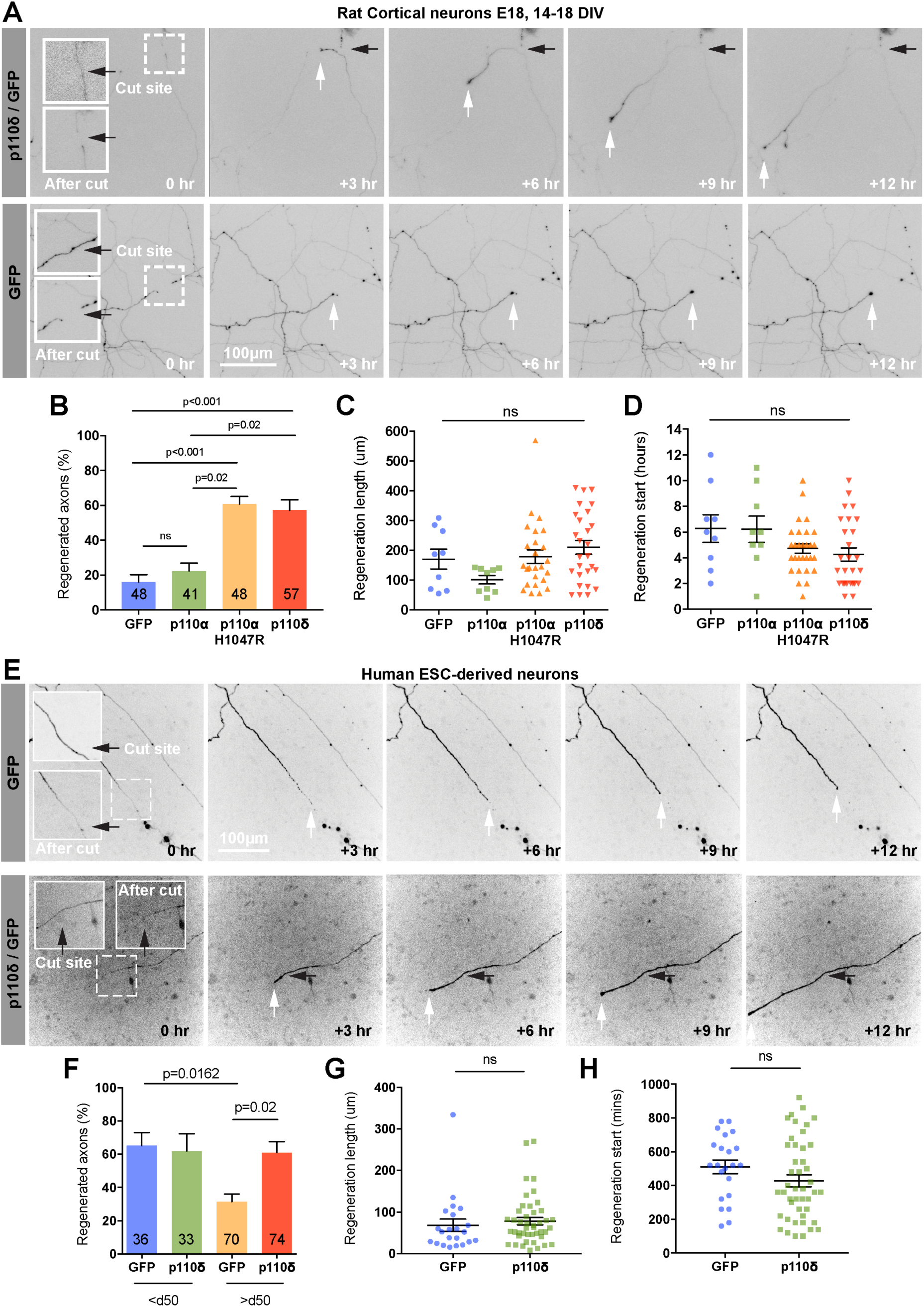
p110δ and p110α^H1047R^ promote axon regeneration of CNS neurons *in vitro*. (A) Axotomised DIV 14-16 cortical neurons expressing p110δ and GFP or GFP alone. Axons were cut >1000 μm from the soma imaged for 14 h. Black arrows mark the cut site, white arrows marks the axon tip. See also supplemental Movies 3 and 4. (B) Percentage of regenerating axons 14 h after laser axotomy. Numbers on bars are injured axons per group. (C) Quantification of axon regeneration length 14 h post-axotomy. (D) Quantification of time to start of regeneration. (E) Axotomised human embryonic stem cell neurons p110δ and GFP or GFP alone. Black arrow marks the cut site, white arrows mark the axon tip. (F) Percentage of regenerating axons of hESC neurons. (G) Quantification of axon regeneration length. Error bars are s.e.m. Data were analysed by Student’s T-test. (H) Quantification of time to start of regeneration. Error bars are s.e.m. Data were analysed by Student’s T-test. (B)-(D), and (F), Error bars are s.e.m. P values indicate significance measured by ANOVA with Tukey’s post-hoc analysis.

We further investigated the translational potential of p110δ by examining axon regeneration in human neurons maturing *in vitro* (human embryonic stem cells (hESC)). We have previously demonstrated that these lose their regenerative ability *in vitro* when cultured beyond 50 days (Koseki et al, 2017). Overexpression of p110δ fully restored the regenerative ability to a level observed for younger neurons (Figure 5E and F). We observed no difference in the length of regenerated axons compared with GFP-expressing control neurons (Figure 5G), but there was a tendency for p110δ expressing neurons to initiate regeneration faster (Figure 5H). Overexpression of p110δ therefore enables CNS axon regeneration in both rat and human neurons, and this is mimicked by hyperactive p110α^H1047R^ whilst overexpression of p110α is ineffective.

### p110δ enhances the axonal transport of regenerative machinery (Rab11 and integrins)

One reason for regenerative failure in CNS neurons is a loss of transport of growth-promoting molecules into axons. Because p110δ expression elevates axonal PIP3 and enhances axon regeneration, we reasoned that this might be accompanied by an increase in the anterograde transport of these receptors. PI3K activity has the potential to trigger anterograde axonal transport through downstream signalling pathways, acting either through the axon transport regulator ARF6 (Eva et al, 2017; Gillingham & Munro, 2007; Macia et al, 2008), or through effects on microtubules via downstream signalling through GSK3β (Mudher et al, 2004). We investigated the axon dynamics of Rab11 and the alpha9 integrin receptor. Enhanced Rab11 transport can overcome the developmental decline in regenerative ability by transporting integrin receptors (Eva et al, 2017). We used spinning disc microscopy to image integrin dynamics in the distal axon by visualising α9-integrin–GFP, in the presence of mCherry (control) or mCherry plus p110δ. The presence of mCherry allows visualisation of the entire axon. Anterograde transport was almost undetectable in control-transfected neurons where we observed predominantly retrograde and static vesicles (Figure 6A and B), in keeping with previous studies (Franssen et al, 2015). Expression of p110δ triggered anterograde movement of integrins, leading to an increase in static integrin vesicles in the distal axon, and an overall increase in the total number of integrin vesicles (Figure 6C). We also analysed the dynamics of Rab11-GFP vesicles, the principle axonal carriers of integrin and other growth receptors (Ascano et al, 2009; Eva et al, 2010; Lazo et al, 2013). Rab11 vesicles moved throughout axons in an oscillatory fashion displaying mostly bidirectional and retrograde movements, in keeping with previous findings (Eva et al, 2017), however expression of p110δ initiated anterograde transport and caused an increase in static and bidirectional movements. This led to an overall increase in Rab11-GFP vesicles in the distal part of p110δ expressing axons, compared with controls (Figure 6 D to E). Expression of p110δ therefore enhances the anterograde transport of alpha9 integrins and Rab11 positive endosomes. These results indicate that increased axonal PIP3 can restore the transport of integrins as well as the Rab11 endosomes that transport them (together with other receptors). Driving these endosomes into axons is sufficient to enable regeneration, indicating enhanced axonal transport as a mechanism by which elevated PI3K activity enables regeneration.

**Figure 6.**
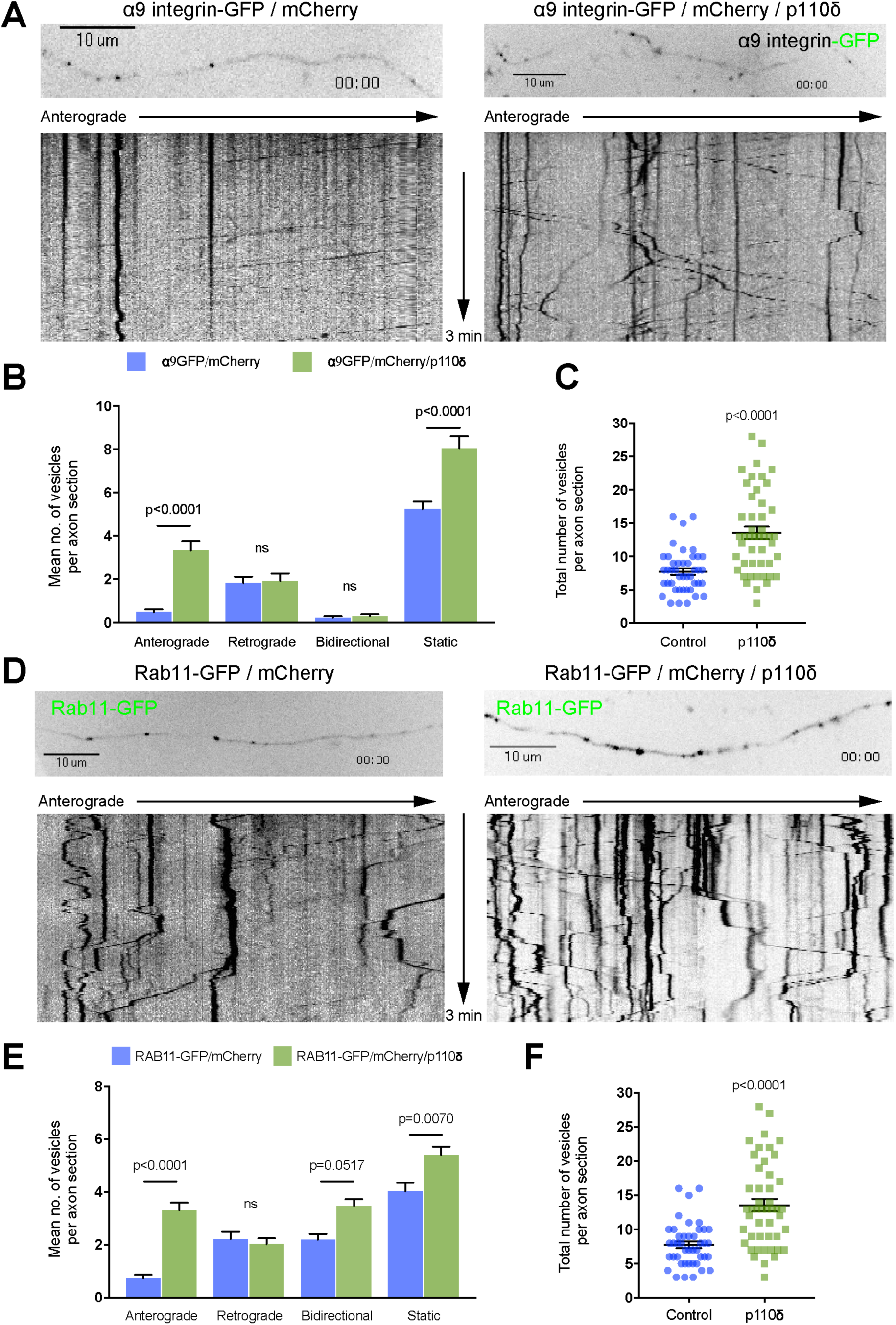
p110δ enhances the axonal transport of integrins and Rab11 endosomes. (A) Kymographs showing dynamics of α9 integrin–GFP in the distal axons of DIV 14-16 neurons, control or co-transfected with p110δ. (B) Quantification of α9 integrin–GFP axon dynamics in the distal axons of DIV 14-16 neurons, control or co-transfected with p110δ. (C) Quantification of total α9 integrin–GFP in the distal axons of DIV 14-16 neurons, control or co-transfected with p110δ. (D) Kymographs showing dynamics of Rab11–GFP in the distal axons of DIV 14-16 neurons, control or co-transfected with p110δ. (E) Quantification of Rab11–GFP axon dynamics in the distal axons of DIV 14-16 neurons, control or co-transfected with p110δ. (F) Quantification of total Rab11–GFP in the distal axons of DIV 14-16 neurons, control or co-transfected with p110δ. (B) and (D) Error bars are s.e.m. P values indicate significance measured by ANOVA with Tukey’s post-hoc analysis. (C) and (F) Error bars are s.e.m. Data were analysed by Student’s T-test.

### Transgenic p110δ and p110α^H1047R^ support RGC survival and axon regeneration in the optic nerve

In order to confirm whether p110δ and p110α^H1047R^ support regeneration in the adult CNS *in vivo*, as we found *in vitro*, we used the optic nerve crush model to examine the effects of PI3K activation on regeneration after a crush injury. We used transgenic mice which conditionally express either p110δ or p110α^H1047R^ from the Rosa26 locus in the presence of Cre recombinase (Figure 7A), and delivered AAV2-Cre-GFP via intravitreal injection. Rosa26 allows expression of a transgene at moderate levels (Nyabi et al, 2009). Two weeks after viral injection, optic nerve crush was performed, and retinal ganglion cell (RGC) survival and optic nerve regeneration were examined 28 days later (Figure 7B). We tested AAV2-Cre-GFP activity using a Cre-reporter mouse which expresses tdTomato from the Rosa26 locus (Figure S3A) and confirmed expression in p110δ and p110α^H1047R^ mice by examining GFP expression (from the viral vector) in the retina (Figure S3B). Activation downstream of PI3K was confirmed by phospho-S6 immunofluorescence. Expression of either p110δ or p110α^H1047R^ led to an increase in the number of cells labelling positive for phospho-S6, however p110α^H1047R^ expression led to a slightly larger increase than p110δ expression (Figure 7C and D). p110δ and p110α^H1047R^ behaved similarly with respect to their effects on RGC survival. The presence of transgenic p110δ or p110α^H1047R^ led to a doubling of the number of cells surviving after 28 days (from 5.5% to 11%), demonstrating a strong neuroprotective effect (Figure 7E and F). We then measured axon regeneration in the optic nerve, and found that p110α^H1047R^ and p110δ again behaved similarly, both enabling a moderate increase in axon regeneration compared with control mice injected with AAV2-Cre.GFP (Figure 7G and H). The results confirm that p110δ and p110α^H1047R^ behave similarly in injured RGC neurons in the CNS *in vivo*, enhancing RGC survival and axon regeneration.

**Figure 7.**
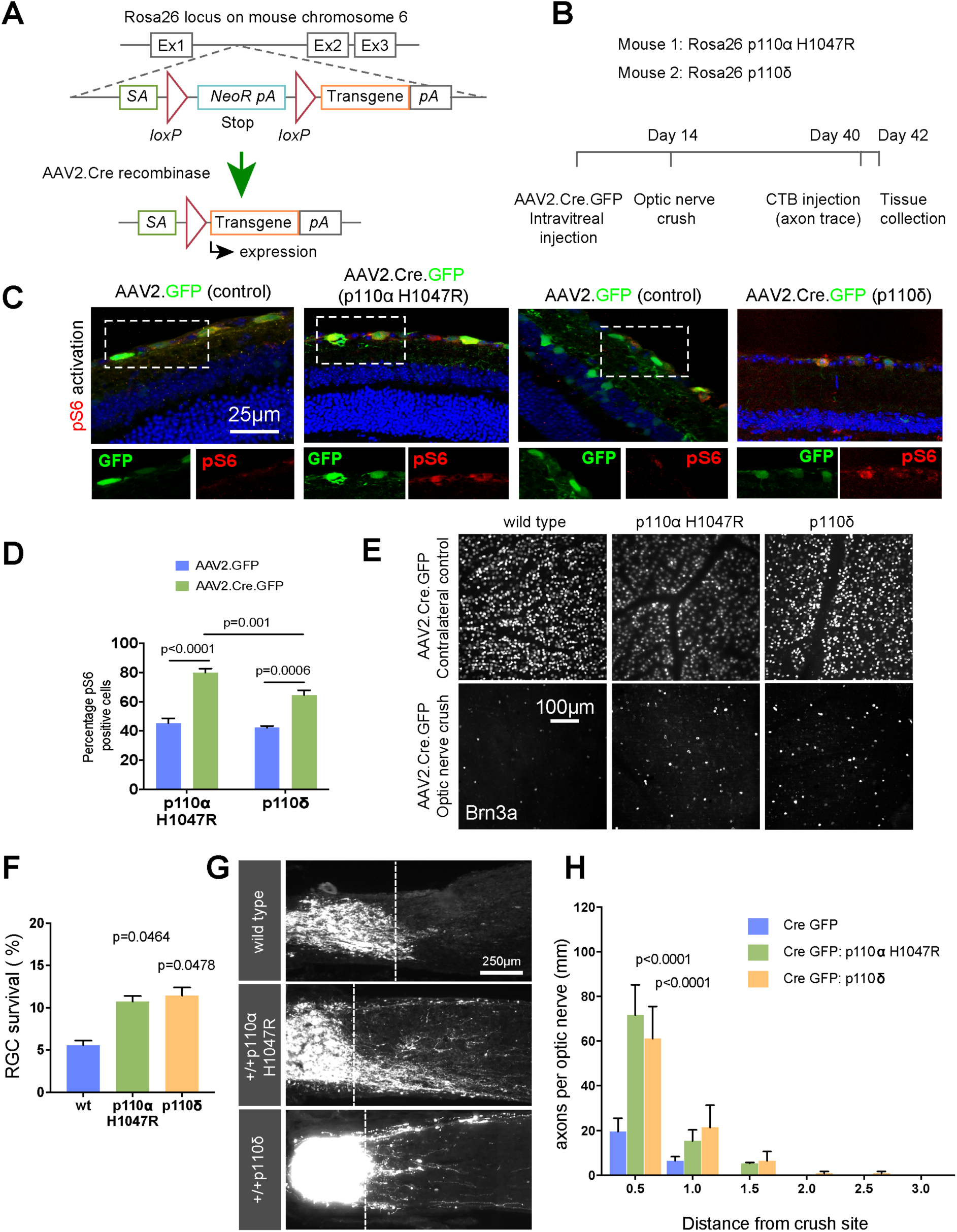
p110δ and p110α^H1047R^ support RGC survival and axon regeneration in the optic nerve. See also supplemental figure S3. (A) Schematic representation of Rosa26 transgene expression via AAV.Cre recombinase. (B) Time course of optic nerve regeneration experiments using Rosa26 p110α^H1047R^ and Rosa26 p110δ mice. C) Retinal sections from AAV-injected mice, immunolabelled for phospho-S6. (D) Percentage of phospho-S6 positive cells in the RGC layer 2 weeks after delivery of AAV-Cre-GFP or AAV-GFP. (E) Retinal whole mounts of AAV-injected mice immunolabelled for the RGC marker Brn3A to indicate cell survival. (F) RGC survival 28 days after optic nerve crush. G) Representative images of CTB-labelled RGC axons 4 weeks after optic nerve crush in wild type (c57bl/6), Rosa26-p110α^H1047R^, and Rosa26-p110δ transgenic mice injected with AAV2.Cre.GFP. (H) Regenerating axons at different distances distal to lesion sites. (D), (F) and (H) Error bars are s.e.m. P values indicate significance as measured by ANOVA with Tukey’s post-hoc analysis.

### AAV2-p110δ facilitates axon regeneration in the optic nerve

The moderate effects of transgenic p110 expression on axon regeneration (described above) were surprising, given the robust effects of p110δ or p110α^H1047R^ expression on CNS axon regeneration *in vitro*. We reasoned that p110 enabled axon regeneration may be dose-dependent, and that the moderate expression generated from the Rosa26 locus (Nyabi et al, 2009) might explain these limited effects. In order to test whether regeneration could be enhanced using a viral gene transfer approach, we produced an AAV2-p110δ construct for viral transduction of RGC neurons via intravitreal injection. We compared this with a similar AAV2-mediated shRNA approach to silence PTEN, the phosphatase responsible for opposing the actions of PI3K by dephosphorylating PIP3 to PIP2. Transgenic suppression of PTEN is another means of stimulating regeneration in the CNS, although virus-mediated shRNA silencing has not proved to be as effective as transgenic deletion (Yungher et al, 2015). We therefore compared viral vector-based delivery of p110δ *versus* viral delivery of a PTEN targeting shRNA. We first confirmed that AAV2-shPTEN-GFP transduction resulted in silencing of PTEN in RGCs compared with AAV2-scrambled-GFP control, PTEN levels being measured by quantitative immunofluorescence (Figure S4A and B). We confirmed transduction of RGCs by AAV2-p110δ immunofluorescence, comparing p110δ with RGC neurons transduced with AAV2-GFP (Figure S4C and D). We examined downstream signalling by labelling for ribosomal phospho-S6. Transduction with AAV2-shPTEN-GFP led to an increase in the percentage of transduced cells labelling positive for phospho-S6 compared with AAV2-scrambled-GFP transduced neurons (from 27% for controls to 48% for PTEN silenced) (Figure 8A and B). Due to the lack of a tag on AAV2-p110δ (due to potential effects on activity), we measured the total number of TUJ1 labelled neurons that also labelled for phospho-S6. Transduction with AAV2-p110δ led to an increase in phospho-S6 positive neurons compared with AAV2-GFP transduced controls (1140 cells for p110δ transduced, compared with 650 cells for GFP transduced) (Figure 8A and C).

**Figure 8.**
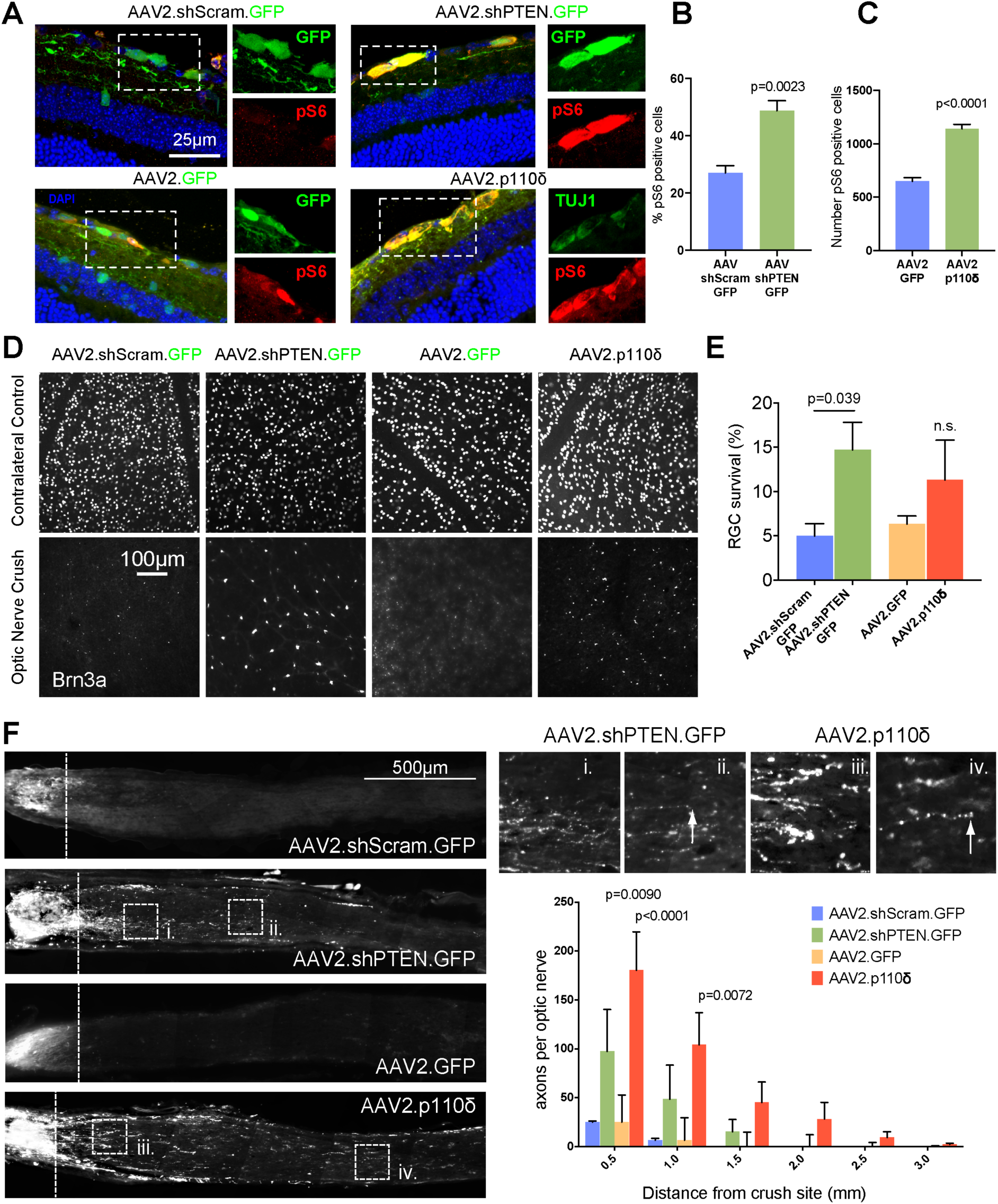
AAV2-p110δ facilitates axon regeneration in the optic nerve. See also supplemental figure S4. (A) Retinal sections from mice injected with AAV2 viruses as indicated, immunolabeled for phospho-S6. (B) Percentage of phospho S6 positive cells in the retinal ganglion layer of mice injected with AAV2.shScramble.GFP or AAV2.shPTEN.GFP. (C) Number of pS6-postive RGCs of mice injected with AAV2.GFP or AAV2.p110δ. (D) Representative images of retinal whole mounts from AAV-injected mice stained for the RGC marker Brn3A to indicate RGC survival. (E) RGC survival 28 days after optic nerve crush. (H) CTB-labelled RGC axons 4 weeks after optic nerve crush. Boxes i. to iv. show regenerating axons. White arrows mark the end of axons. Graph shows quantification of regenerating axons at different distances distal to the lesion sites. (B) and (C) Error bars are s.e.m. P values indicate significance measured by Students T-test. (E) and (H) Error bars are s.e.m. P values indicate significance measured by ANOVA with Tukey’s post-hoc analysis.

Having validated the two approaches, we next compared their effects on RGC survival and axon regeneration after optic nerve crush. Silencing of PTEN led to a robust increase in the survival of RGC neurons compared with controls at 28 days after crush (14.7% of PTEN silenced neurons survived compared to 4.9% of control neurons), whilst transduction with p110δ led to a smaller effect on survival (11.3% of p110δ transduced neurons compared 6.3% of GFP control neurons) (Figure 8D and E), similar to the amount observed due to transgenic expression (Figure 7F). We examined CTB-traced RGC axon regeneration in the optic nerve, and found that both AAV2 mediated silencing of PTEN and AAV2-delivery of p110δ enhanced axon regeneration. Transduction with p110δ had the most robust effects on regeneration, with 180 axons counted at 0.5 mm into the optic nerve, compared with 25 axons in control-injected mice (Figure 8F). Injection of AAV2-shPTEN-GFP led to 97 axons at 0.5 mm, again compared with 25 axons for controls (this compares with 60 axons for p110δ, 70 for p110α^H1047R^ with transgenic expression). AAV2-p110δ also enabled axons to regenerate further into the optic nerve reaching a maximum distance of 3 mm, whilst silencing of PTEN enabled axons to reach a maximum of 1.5 mm (transgenic expression of p110α^H1047R^ and p110δ also gave 1.5 mm regeneration). These data demonstrate that the PI3K pathway can be targeted to stimulate RGC survival and axon regeneration either by expressing p110δ or by silencing PTEN, but that expression of p110δ has the most robust effects on axon regeneration. Together these data show that enhancing PI3K activity in CNS neurons greatly enhances their ability to regenerate their axons, and indicate viral delivery of p110δ to CNS neurons as a novel approach to boost signalling through this regenerative pathway.

## Discussion

Our study aimed to find a new method of stimulating axon regeneration based on manipulation of PIP3 levels in neurons. In previous work the approach to this has mainly been to knock down the PIP3 dephosphorylating enzyme PTEN (Geoffroy et al, 2015; Lee et al, 2014; Liu et al, 2010; Park et al, 2008). However, this approach depends on generation of PIP3 by PI3K, and if there is little PIP3 being produced, PTEN knockdown will have little effect. We reasoned that limited PIP3 production in mature neurons would explain the observation that PTEN knockout is effective at promoting axon regeneration in immature neurons, less so in the fully adult CNS. We used a new method to visualise PIP3 levels in PNS and CNS neurons at different levels of maturity. In sensory axons, which regenerate readily, we observed high levels of PIP3 in the growth cones, dynamically changing with growth cone movements. In cortical CNS neurons, developing immature neurons exhibited intense PIP3 levels in their axon and growth cone during the period of rapid axon growth. However, PIP3 levels decreased to a much lower level at a time when transport declines and axons lose their capacity for regeneration. We also investigated the role of PI3K isoforms in regeneration in PNS neurons using isoform-specific inhibitors. In sensory axons, which regenerate successfully and quickly, growing in microfluidic chambers to separate axonal and cell body compartments, inhibitors of p110α blocked both growth and post-axotomy regeneration when applied to either compartment, but a p110δ inhibitor blocked regeneration without affecting normal growth. This supports a previous study that links p110δ to sensory regeneration (Eickholt et al, 2007) and suggests that p110δ is uniquely linked to regeneration. Overexpression of p110 isoforms was therefore tested for the ability to increase PIP3 levels and to promote regeneration. Overexpression of p110δ in mature CNS neurons partially restored PIP3 levels, particularly in axon tips but p110α was ineffective. However, introducing the hyperactivating H1047R mutation to p110α had the same effect on PIP3 levels as p110δ.

Stimulation of axon regeneration was therefore assessed. p110δ and p110α^H1047R^ transfected into mature cortical neurons strongly enhanced axon regeneration. In the CNS *in vivo* transgenic expression led to enhanced neuroprotection in the retina, and regeneration in the optic nerve after a crush injury. Importantly, viral transduction of p110δ (which produces a higher level of expression) into adult RGC neurons led to axons regenerating for a greater distance after injury, demonstrating a novel approach to boost CNS regeneration through the PI3K pathway by gene transfer. Previous work has indicated that the hyperactive H1047R mutation of p110α can behave like p110δ to sustain AKT signalling in fibroblasts, by functioning independently of co-activation by Ras (Kang et al, 2006). We found that expression of native p110α had no effect on PIP3 generation, it was only p110δ or p110α^H1047R^ that enhanced axonal PIP3 (Figure 3). This suggests that receptor activation of PI3K is at a low level in mature CNS neurons, and that p110δ can potentiate PI3K signalling in low trophic conditions that are insufficient to activate native p110α. Our findings suggest that p110δ has a lower threshold of activation and that signals normally required to fully activate p110α (such as adhesion and growth factor receptors and Ras) may not be available in the axon. This has important implications for understanding the nature of p110δ beyond the nervous system, and may help to explain how T-cells utilise p110δ for development, differentiation and function (Okkenhaug, 2013).

The PI3K signalling pathway is a well-known regulator of axon regeneration, based on seminal studies which demonstrated that deletion of PTEN leads to robust regeneration in the CNS through downstream signalling via mTOR (Park et al, 2008). Aside from mTOR, the mechanism through which PTEN deletion stimulates regeneration is not completely understood. One difficulty in understanding its mechanism is that in addition to functioning as a lipid phosphatase (PTEN opposes PI3K by converting PIP3 back to PIP2) it also has protein phosphatase activity (Kreis et al, 2014). It is now also apparent that PTEN not only dephosphorylates PI(3,4,5)P_3_, but also functions as a PI(3,4)P_2_ phosphatase, a role linked with cancer invasion (Malek et al, 2017). Our findings argue in favour of PTEN functioning through the regulation of PIP3, and confirm the importance of this molecule in the regulation of axon regeneration.

Most PI3K signalling events rely on more than one isoform (Hawkins & Stephens, 2015), and whilst the p110δ isoform contributes to sciatic nerve regeneration (Eickholt et al, 2007), the contribution of the other isoforms remained unknown. Our findings demonstrate that both p110α and δ are required for efficient axon regeneration, and that p110α functions in both the axon and cell body, whilst p110δ is specifically required in the axon (Figure 1). Inhibition of p110α opposed both axon growth and regeneration, whilst the action of p110δ was specific to regeneration, with the p110δ inhibitor blocking regeneration but not growth of uncut axons. Taken together, these results suggest that p110α mediates the somatic and axonal signalling that is necessary to support adult DRG axon growth (itself a regenerative phenomenon), whilst p110δ is further required within the axon to facilitate the redevelopment of a growth cone after injury. In CNS neurons, overexpression of either p110δ or p110α^H1047R^ was sufficient to enable efficient regeneration both *in vitro* and *in vivo,* and AAV mediated p110δ expression in RGCs promoted robust optic nerve regeneration.

Our findings demonstrate the importance of PIP3 in the axon as well the cell body for optimal regeneration. Studies targeting the PTEN/PI3K pathway for CNS axon regeneration examined downstream activation by measuring ribosomal S6 phosphorylation in the cell body. An important action of the observed increase in axonal PIP3 caused by expression of p110δ was to restore the anterograde transport of integrins and the Rab11 endosomes that transport integrin and other receptors into mature CNS axons. Previous work has shown that forcing Rab11 vesicles into these mature axons promotes regeneration, so the restoration of transport is one mechanism through which p110δ transfection can restore regeneration. This suggests a feed-forward mechanism because many of these receptors are also activators of PI3K. The exclusion of PI3K-activating receptors from mature axons is probably a reason why the generation of PIP3 is low in mature axons, and why overexpression of p110α in neurons did not increase PIP3 levels. PTEN deletion and presumably p110δ expression also enhance regeneration via mTOR signalling in the cell body, however mTOR has recently been found to be present at the growth cone of developing axons in the mouse cerebral cortex, suggesting the PI3K-AKT-mTOR pathway may also function locally within the axon (Poulopoulos et al, 2019). It is important however to note that signalling through mTOR only represents a subset of the signalling molecules regulated downstream of PIP3 or PIP2. PIP3 and PIP2 exert their effects via a wide variety of proteins with PH domains that may also contribute to regenerative effects, including the regulatory molecules of small GTPases that regulate the cytoskeleton, such Rac1 and Cdc42 (Sosa et al, 2006; Welch et al, 2002; Yoshizawa et al, 2005), or that regulate axon transport such as ARF6 and Rab11 (Gillingham & Munro, 2007; Nieuwenhuis & Eva, 2018). Enhancing PI3K signalling may therefore promote the axonal transport and trafficking of a range of axon growth-promoting molecules that are normally excluded (Eva et al, 2017; Hollis et al, 2009a; Hollis et al, 2009b). The importance of targeting the axon as well as the cell body is often overlooked, however there is increased axonal transport as part of the PNS injury response and in the CNS, increased transport enables regeneration (Petrova & Eva, 2018).

The search for translatable methods for promoting regeneration in the damaged CNS continues. A successful strategy will likely involve multiple interventions. Could manipulation of PI3K be translationally useful? PTEN deletion or p110α^H1047R^ expression can be oncogenic in dividing cells. However, expression under a neuron-specific promoter targets expression to a non-dividing cell type, and it is unlikely that expression of p110δ in CNS neurons would lead to cell transformation, particularly given its usual expression in PNS neurons. Our study puts forward AAV-mediated delivery of p110δ as a novel means of stimulating regeneration through the PI3K pathway in a potentially translatable fashion. We propose that p110δ should also be considered as an additional intervention with other regenerative strategies that target the PI3K pathway, either through growth factor treatments such as IGF-1 plus osteopotin, (Duan et al, 2015; Liu et al, 2017), expression of activated integrins (Cheah et al, 2016) or pharmacological interventions such as insulin (Agostinone et al, 2018).

## Supporting information

Barber et al Movie 1

Barber et al Movie 2

Barber et al Movie 3

Barber et al Movie 4

## Acknowledgements

We acknowledge the Babraham Institute Gene Targeting facility for the generation of the Rosa26 p110 mouse strains, and would like to thank Dr. Len Stephens and Dr. Phill Hawkins (Babraham) for the gift of the GFP-AKT-PH mouse strain, and for their advice and support. The work was supported by grants from MRC-Sackler, International Spinal Research Trust (NRB110), Wellcome Trust, ERA-NET NEURON grant AxonRepair (013-16-002), Medical Research Council MRC (MR/R004544/1, MR/R004463/1), Christopher and Dana Reeve Foundation, Cambridge Eye Trust, Fight for Sight, NERC and a core support grant from the Wellcome Trust and MRC to the Wellcome Trust – Medical Research Council Cambridge Stem Cell Institute. B.J.E. funding was provided by the DFG (SFB 958, TP16; SFB TRR 186, TP10). S.v.E was funded by EMBO ALTF 1436-2015, and S.v.E & C.ff-C by MS Society UK. K.O. laboratory funding was provided by the BBSRC and the Wellcome Trust.

## Author contributions

A.C.B, R.S.E, B.N., C.S.P, J.F, P.D.S, S.v.E and R.E. performed and analysed experiments. A.R.M generated PI3K transgenic mice. B.H., A.O. and L.A.H. generated AAV.p110delta. R.C. J.C. and S.S.D. analysed RGC survival. K.O. designed and supervised transgenic generation of mice. B.J.E designed and supervised PIP3 detection experiments. R.E., J.W.F, K.R.M, A.C.B, C.ff-C, S.v.E, B.J.E and K.O. conceived designed and supervised experiments and obtained funding. R.E., B.N., A.C.B. and R.S.E prepared movies and figures. R.E. wrote the manuscript.

## Materials and Methods

### Mouse Strains

C57BL6/J mice were used during this study, as well as four transgenic mouse strains: GFP-AKT-PH, Rosa26 p110α^H1047R^, Rosa26 p110δ and B6;129S6-Gt(ROSA)26Sortm14(CAG-tdTomato)Hze/J (https://www.jax.org/strain/007908). These are described in detail in the supporting information section.

### DRG culture

Dissociated DRG neuronal cultures were obtained from adult male Sprague Dawley (SD) rats and from the transgenic AKT-PH-GFP adult mouse. DRGs were incubated with 0.1% collagenase in Dulbecco’s modified Eagle’s medium (DMEM) for 90 min at 37°C followed by 10 min in trypsin at 37°C, and dissociated by trituration. Dissociated cells were centrifuged through a layer of 15% bovine serum albumin (BSA) and cultured on 1 µg/ml laminin on glass-bottom dishes (Greiner) in DMEM supplemented with 10% fetal calf serum, and 50 ng/ml nerve growth factor (NGF). For compartmentalised experiments, DRG neurons were plated in Xona microfluidic devices (Xona SND150) on glass coverslips. For PI3K inhibitor experiments, media was exchanged for serum free media overnight. Separation of media was achieved by maintaining a pressure gradient between the axonal and somatic side of the device.

### Embryonic cortical neuron culture

Primary rat cortical neuron culture has been described previously (Eva et al, 2017). Cultures were prepared from embryonic day 18 (E18) SD rats. Neurons were dissociated with papain for 8 min at 37°C, washed with HBSS and cultured in MACS Neuromedium supplemented with MACS Neurobrew (Miltenyi), and plated on glass-bottom dishes (Greiner) coated with poly-D-lysine. Cells were transfected at 10 DIV, and experiments performed between DIV 14 and 17. Cortical neurons were transfected with magnetic nano-particles (Franssen et al, 2015).

### hESC dopaminergic neuron culture

RC17 hESC cell culture has been have been previously described in detail before (De Sousa et al, 2016; Koseki et al, 2017). Cells (RRID:CVCL_L206) were sourced from Roslin Cells, Scottish Centre for Regenerative Medicine, Edinburgh, UK. The cell line is free from mycoplasma contamination as determined by RT-qPCR. On d0 hESC were detached and transferred to form embryoid bodies from d0 to d4 in neural induction medium (Neurobasal:DMEM/F12 (1:1), 0.2% P/S, L-glutamine, N2, B27, recombinant Sonic Hedgehog Shh (C24II), recombinant noggin (Ng), SB431542 (SB) and CHIR99021 (CH). Rock inhibitor (RI,) was present in the medium from d0 to d2. At d4, embryoid bodies were plated in poly-L-ornithine Laminin and Fibronectin (PLF) coated plates and cultured in neural proliferation medium (Neurobasal:DMEM/F12 (1:1), 0.5xN2, 0.5xB27, supplemented with Shh, Ng, SB and CH from d4-d7, and with Shh, Ng and CH from d7-d9). At d11, cells were dissociated with Accutase and 50*104 cells per well single cell suspensions were plated in PLF-coated 8-well chamber slides. Cells were cultured in neuronal differentiation medium (NB with 0.2% P/S, l-glutamine, B27, Ascorbic Acid, recombinant human BDNF, GDNF, db-CAMP; 50 mM) and DAPT from d14 onwards. Medium was replaced twice weekly up to day 50, after which medium was replaced weekly.

### N1E Cell Culture

N1E-115 cells (ATCC) were plated at a density of 5.000 cells / cm² in DMEM-high glucose with GlutaMAX (Gibco) + 10% FBS and incubated for 6-7 h at 5% CO_2_ and 37°C.

### DNA Constructs

Constructs for expressing p110 were generated from pHRsinUbEm, a bicistronic vector expressing EGFP under the control of the SFFV promoter and emerald fluorescent protein under the control of the Ubiquitin promoter. p110δ (PIK3CD) was a gift from Roger Williams (MRC Laboratory for Molecular Biology, UK). PIK3CD was cloned from pcDNA3.1 in place of EGFP. p110α (PIK3CA) or p110α H1047R were cloned into the same site from pBabe puro vectors. pBabe puro HA PIK3CA and p110α H1047R were gifts from Jean Zhao (Addgene plasmid #12522 and #12524; http://n2t.net/addgene:12522; RRID:Addgene_12522, http://n2t.net/addgene:12524; RRID:Addgene_12524) (Zhao et al, 2005). AAV.CAG.p110δ was generated by cloning the human PIK3CD sequence into an AAV-sCAG vector backbone by Gibson assembly (Gibson et al, 2009).

### Antibodies

PI(3,4,5)P: anti-PtdIns(3,4,5)P_3_ monoclonal antibody (Z-P345b, 1:200, Echelon Biosciences), GFP (rabbit, 1:500 Abcam ab290), phospho-S6 ribosomal protein (Ser235/236) (91B2) (rabbit, 1:200, Cell Signaling 4857S), TUJ1(βIII Tubulin) (mouse, 1:400, Promega G7121), PTEN (D4.3) XP (rabbit, 1:100, Cell Signalling 9188S), p110δ (rabbit, 1:500, Abcam ab1678), Brn3a (C-20) (goat, 1:200, Santa Cruz sc-31984). Anti-goat Alexa Fluor-647 (1:1000, A21447, Life technologies). Anti-rabbit IgG conjugated Alexa Fluor 568 (A10042, 1:1000, ThermoFischer scientific). Anti-mouse IgG conjugated Alexa Fluor 568 (A11004, 1:1000, ThermoFischer scientific). Anti-rabbit IgG conjugated Alexa Fluor 488 (A-21206, 1:1000, ThermoFischer scientific). Anti-mouse IgG conjugated Alexa Fluor 488 (A-21202, 1:1000, ThermoFischer scientific).

### Small molecule inhibitors

The following small molecule inhibitors were used to inhibit the various isoforms of p110. The indicated concentrations were chosen based on their known IC50, and from previously reported cell culture experiments: Pan-p110 (p110α/β/δ): LY294002 20 μM (IC50 (in cell free assays) 0.5 μM/0.97 μM/0.57 μM, respectively). p110α: A66 5 μM (IC50 32nM). p110α/δ: XL-147 5 μM (IC50 39 nM/ 36nM). p110β: TGX221 500 nM (IC50 5 nM). p110δ: IC-87114 10μM (IC50 0.5 μM). p110δ: Idelalisib 500 nM (IC50 2.5 nM).

### Virus production and injection

Three viruses were sourced commercially: AAV2.CMV.Cre.GFP (Vector Biolabs, Catalog #7016), AAV2.CMV.GFP (Vigene Biosciences, Catalog #CV10004) and AAV2.U6.shRNA(scramble).CMV.GFP (SignaGen Labs, Catalog #SL100815). AAV2.U6.shPTEN.CMV.GFP, was a gift from Zhigang He, (Boston Children’s Hospital) and AAV2.CAG.p110delta was produced by Vigene Biosciences.

Mice received 2 μl intravitreal injections of AAV. All viruses were injected into the left eye only at 1 x 10^13^ GC/ml. For validation experiments, mice also received intravitreal injection of the appropriate control into the right eye.

### Optic nerve injury

Animal work was carried out in accordance with the UK Home Office regulations for the care and use of laboratory animals, the UK Animals (Scientific Procedures) Act (1986), and the Association for Research in Vision and Ophthalmology’s Statement for the Use of Animals in Ophthalmic and Visual Research. Surgical procedures were performed under anesthesia using intraperitoneal injection of ketamine (100 mg/kg) and Xylazine (10 mg/kg).

Optic nerve injuries were carried out as previously described (Smith et al, 2009). The optic nerve behind the left eye was exposed intraorbitally, and crushed with fine forceps for 10 sec, approximately 0.5 mm behind the optic disc. 26 days after the injury, mice received a 2 µl intravitreal injection of cholera toxin subunit β (CTB) with an Alexa Fluor 555 conjugate at 1mg/ml. 28 days post crush animals were perfused with 4% paraformaldehyde (PFA) and the eyes and optic nerves collected for analysis.

### Immunohistochemistry

Retinas were fixed in 4% PFA for 2 h. Whole-mounts were washed four times with 0.5% PBS-TritonX100. In between the second and third wash, a permeation step was performed to improve antibody penetration by freezing the retinas in 0.5% PBS-TritonX100 for 10 min at −70°C, and washing was continued after thawing. Optic nerves were fixed overnight at 4°C in 4% PFA, followed by 30% sucrose overnight at 4°C. Sections were blocked with 2% BSA, 10% donkey serum in 2% PBS-TritonX100. Primary antibodies were incubated at 4°C overnight, secondary antibodies for 2 h at room temperature.

### Standard immunocytochemistry

Cortical neurons were fixed with 3% paraformaldehyde (PFA) in PBS for 15 min and permeabilized with 0.1% Triton X-100 in PBS for 5 min. Cells were blocked with 3% bovine serum albumin (BSA) in PBS for 1 h. After blocking, the cells were incubated with primary antibodies diluted in 3% BSA in PBS at 4°C overnight. Secondary antibodies that were diluted in 3% BSA in PBS for 1 h at room temperature.

### Phospholipid fixation and immunocytochemistry

We adapted a fixation technique previously used to detect PI(4,)P_2_ and PI(4,5)P_2_ (Hammond et al, 2009). Cortical neuron cultures were fixed using pre-warmed (37°C) 3% formaldehyde and 0.2% glutaraldehyde (GA; G011/3, TAAB Laboratories) in PBS for 15 min at room temperature, washed in 50 mM NH_4_Cl in PBS, then maintained at 4°C. Cells were incubated with 4°C blocking and permeabilisation solution (0.2% saponin, 50 mM NH_4_Cl and 3% BSA in PBS) for 30 min, then with anti-PtdIns(3,4,5)P3 (Z-P345b, 1:200, Echelon Biosciences) in blocking solution for 3 h before washing three times in 50 mM NH_4_Cl 30 min. Secondary antibody was applied for 2 h at 4°C. After another 30 min wash, cells were post-fixed in 3% formaldehyde in PBS for 5 min at 4°C before being moved to room temperature for a further 10 min.

### Insulin Stimulation of N1E cells

Cells were starved in DMEM without serum overnight in the presence of either 500 nM GDC-0941 (Selleckchem) or an equal volume of cell-culture grade DMSO (Applichem). Cells were stimulated with 20 µg/ml insulin (Sigma) in DMEM or control treated with equal amounts of DMEM for 1 min prior to fixation in PIP3-Immobilization fixative and stained with for PIP3 (anti-PIP3, Echelon) and F-actin (Phalloidin, Thermo Fisher Scientific).

### Confocal and widefield microscopy

Laser-scanning confocal microscopy was performed using a Leica DMI4000B microscope, and a Leica TCS SPE confocal system controlled with Leica LAS AF. Fluorescence and wide-field microscopy were performed using a Leica DMI6000B with a Leica DFC350 FX CCD camera and a Leica AF7000 with a Hamamatsu EM CCD C9100 camera and Leica LAS AF. Leica AF7000 was used for imaging of axon and growth cone regeneration after axotomy.

### TIRF Microscopy

Total internal reflection fluorescence (TIRF) microscopy was carried out on a Leica DMI6000B adapted with a dedicated TIRF module from Rapp OptoElectronic. GFP was excited with a 488 nm laser and images were acquired using a Leica Plan Apo 100x / NA1.47 Oil TIRF objective and Hamamatsu EM CCD C9100 camera controlled by Leica LAS AF and Rapp OptoElectronic software.

### Laser axotomy of DRG neurons

Laser axotomy of DRG neurons was performed as described previously (Eva et al, 2017). Axons were cut directly before a growth cone, to determine the proportion of axons that regenerate rapidly after injury. Cultures were serum starved overnight, and inhibitors were added at the start of the experiment. Axons were severed using a 355 nm DPSL laser (Rapp OptoElectronic, Hamburg, Germany) connected to a Leica DMI6000B, and images were acquired every 15 min for 2 h. Successful regeneration was determined as the development and extension of a new growth cone.

### Laser axotomy of cortical neurons

Laser axotomy of cortical neurons was performed as described previously (Eva et al, 2017; Koseki et al, 2017) Axons were severed using a 355 nm DPSL laser (Rapp OptoElectronic, Hamburg, Germany) connected to a Leica DMI6000B. Cortical neurons were axotomised at DIV 14–17 at distances of 800–2000 µm from the cell body on a section of axon free from branches. Images after axotomy were acquired every 30 min for 14 h. Regeneration was classed as the development of a new growth cone followed by axon extension for a minimum of 50 µm.

### Laser axotomy of hESC-derived neurons

Laser axotomy of hESC-derived neurons was performed as described previously (Koseki et al, 2017). Axons were severed using a 365 nm laser (Micropoint, Andor) connected to an Andor spinning disk confocal microscope. Neurons were axotomised at d45-65 at distances of >500µm distal to the cell body on a section of axon free from branches. A single axon cut was made per neuron. Images after axotomy were acquired every 20 min for 16 h. Regeneration was classed as the development of a new growth cone followed by axon extension for a minimum of 50 µm.

### Quantification of PIP3 in cortical neurons

E18 cortical neurons were fixed at DIV 3, 8, or 16 and PIP3 was detected as described above. All cultures were fixed and labelled using identical conditions. Images were acquired by confocal laser-scanning microscopy using a Leica TCS SPE confocal microscope. Identical settings were used for each image using Leica LAS AF. Z-stacks were acquired for each image, spanning the entire depth of either the cell body or the growth cone. PIP3 fluorescence intensity was measured using Leica LAS AF. Immunofluorescence intensities were calculated by measuring the region of interest (ROI), and subtracting the intensity of a control region adjacent to the ROI.

### Quantification of PIP3 in N1E cells

Imaging was performed on a Leica SP5 confocal system with a 40x Objective with identical settings across experiments. Stacks covered the complete height of all cells with. Areas with comparable cell densities were selected in the F-actin channel. Mean PIP3 intensity in N1E cells was quantified in Fiji (ImageJ 1.51n) on max-projections. Intensities were compared in 15 images per treatment from 3 independent cultures.

### Analysis of neuronal morphology

Images were captured on a Leica DMI6000B, with a 40X-oil objective using Leica LAS AF. Semi-automated and standardized analysis was performed using MATLAB platform version 2017 and SynD (Schmitz et al, 2011). The output of SynD was used for data analysis of dendritic length, fluorescent intensities, sholl analysis, and soma size.

### Neurite outgrowth assay

Cultured cortical neurons were transfected at 2 DIV and fixed at 4 DIV. Images for neuronal morphology were acquired using a Leica DMI6000B microscope, with a 40X-oil objective using Leica LAS AF. Neurite lengths were measured using the ImageJ plugin ‘simple neurite tracer’(Longair et al, 2011).

### Axon transport analysis

Axon transport analysis of integrins and Rab11 has been described in detail before (Eva et al, 2017; Franssen et al, 2015). Briefly, Cortical neurons were transfected at DIV 10 with α9 integrin–GFP, or Rab11–GFP together with either mCherry (control) or mCherry plus p110δ, and imaged at DIV 14–16 DIV using spinning disc confocal microscopy. Sections of axons were imaged at a region in the distal part of the axon (>800 µm from the cell body). Vesicles were tracked for their visible lifetime, and analysed by kymography to classify the proportion of vesicles classed as anterograde, retrograde, bidirectional or immobile per axon section. Vesicles with a total movement less than 2 µm during their visible lifetimes were classed as immobile. Vesicles moving in both directions but with net movement of less than 2 µm (during their visible lifetimes) were classed as bidirectional. Vesicles with net movements greater than 5 µm in either direction by the end of their visible lifetimes were classed as anterograde or retrograde accordingly.

### Optic nerve regenerating axon counts

To measure regenerating RGC axons after optic nerve crush, longitudinal sections of optic nerves were serially collected. Regenerating RGC axons were quantified as described previously (Smith et al, 2009), by counting the number of CTB labelled axons at the indicated distances beyond the crush site from 4 sections per optic nerve. Axonal sections were imaged using a Zeiss AxioScan Z1 at x40 magnification.

### RGC survival counts

For retinal whole-mounts, 2 images were taken from each of the four retinal quadrants at x20 magnification, sampling both the more central and the more peripheral region of each quadrant. Images were then analysed in Image J Fiji using Image-Based Tool for Counting Nuclei (ITCN) Plugin (University of California, Santa Barbara, CA, USA) to count Brn3A-labelled cells. The number of RGCs in the left injured eye was expressed as a percentage survival value compared to the mean number of RGCs of the contralateral control eyes.

### RGC fluorescence analysis

To confirm successful viral transduction, eye cup images were stained for GFP or PI3KDelta counterstained with DAPI, and imaged using a Zeiss AxioScan Z1 at x20 magnification. Retinal sections immunolabelled for pS6 were examined by fluorescence microscopy. 100 GFP-positive RGCs were counted and identified as positive or negative for pS6. For AAV2.p110δ transduced RGCs, the total number of pS6 positive cells was counted from 12 retinal sections throughout the eye compared to total number of pS6 positive cells from the control group using TUJ1 to identify neurons. Fluorescence intensity was measured using ImageJ FIJI.

### Quantitative and Statistical Analysis

Statistical analysis was performed throughout using Graphpad Prism. Fisher’s exact test was calculated using Graphpad online (https://www.graphpad.com/quickcalcs/contingency1.cfm). Data were analysed by ANOVA with post hoc analysis, Student’s t-test and Fisher’s exact test. Sample sizes were based on previously published data using similar techniques. Statistical analysis was performed using ANOVA followed by Tukey’s multiple comparison test or Student’s T-test, as indicated in the figure legends. Percentage of regenerating axons was compared by Fisher’s exact test.

