## Supplemental Data 1 for "PI 3-kinase delta enhances axonal PIP_3_ to support axon regeneration in the adult CNS"

### Supplemental Information

### Supplemental Information

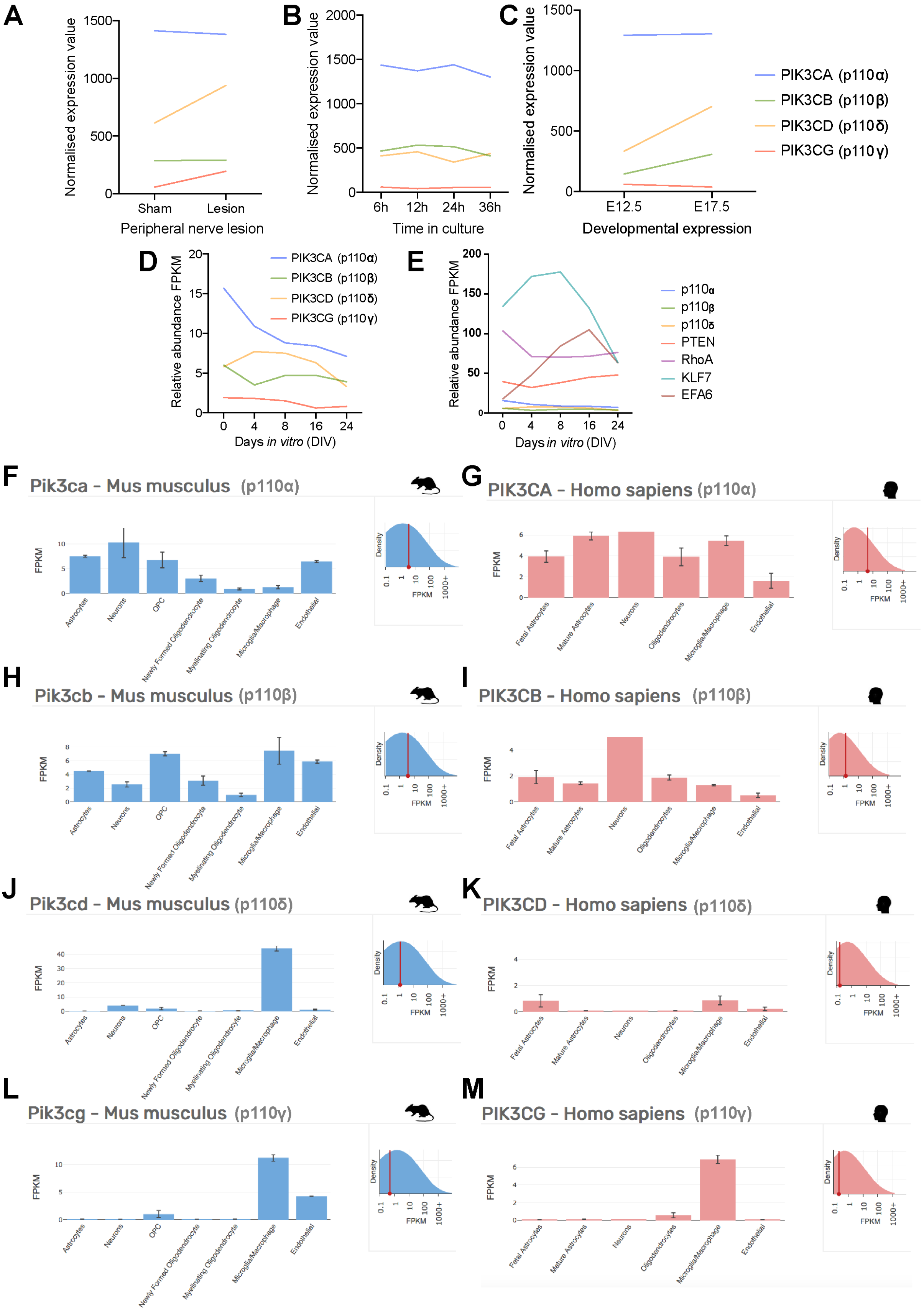


**Figure S1.**

**Figure S1. Gene expression profile of p110 isoforms in the nervous system from previously published RNAseq datasets.**

Panels A to C show data from Tedeschi et al 2016, Panels D and E from Koseki et al 2017, and Panels F to M from Brain-RNAseq databases (http://www.brainrnaseq.org/), Zhang et al 2014 and 2106.

(A) Normalized mean expression values of p110 genes in adult mouse DRG neurons isolated after sciatic nerve lesion compared to a sham control.

(B) Normalized mean expression values of p110 genes in cultured mouse DRG neurons 6-, 12, 24-, and 36 h post-plating (representing the shift from arborizing to elongating axon growth).

(C) Normalized mean expression values of p110 genes in cultured mouse DRG neurons from at embryonic day 12.5 and 17.5.

(D) Relative abundance of p110 mRNA levels in cortical neurons cultured from E18 rat embryos at increasing periods of time *in vitro*.

(E) Relative abundance of p110 mRNA levels in cortical neurons cultured from E18 rat embryos at increasing periods of time *in vitro*, also showing expression levels other regeneration-associated genes.

(F - M) Relative abundance (FPKM) of p110 genes in various mouse and human brain cell types (astrocytes, neurons oligodendrocyte precursor cells, newly formed oligodendrocytes, myelinating oligodendrocytes, microglia/macrophages, and endothelial cells).

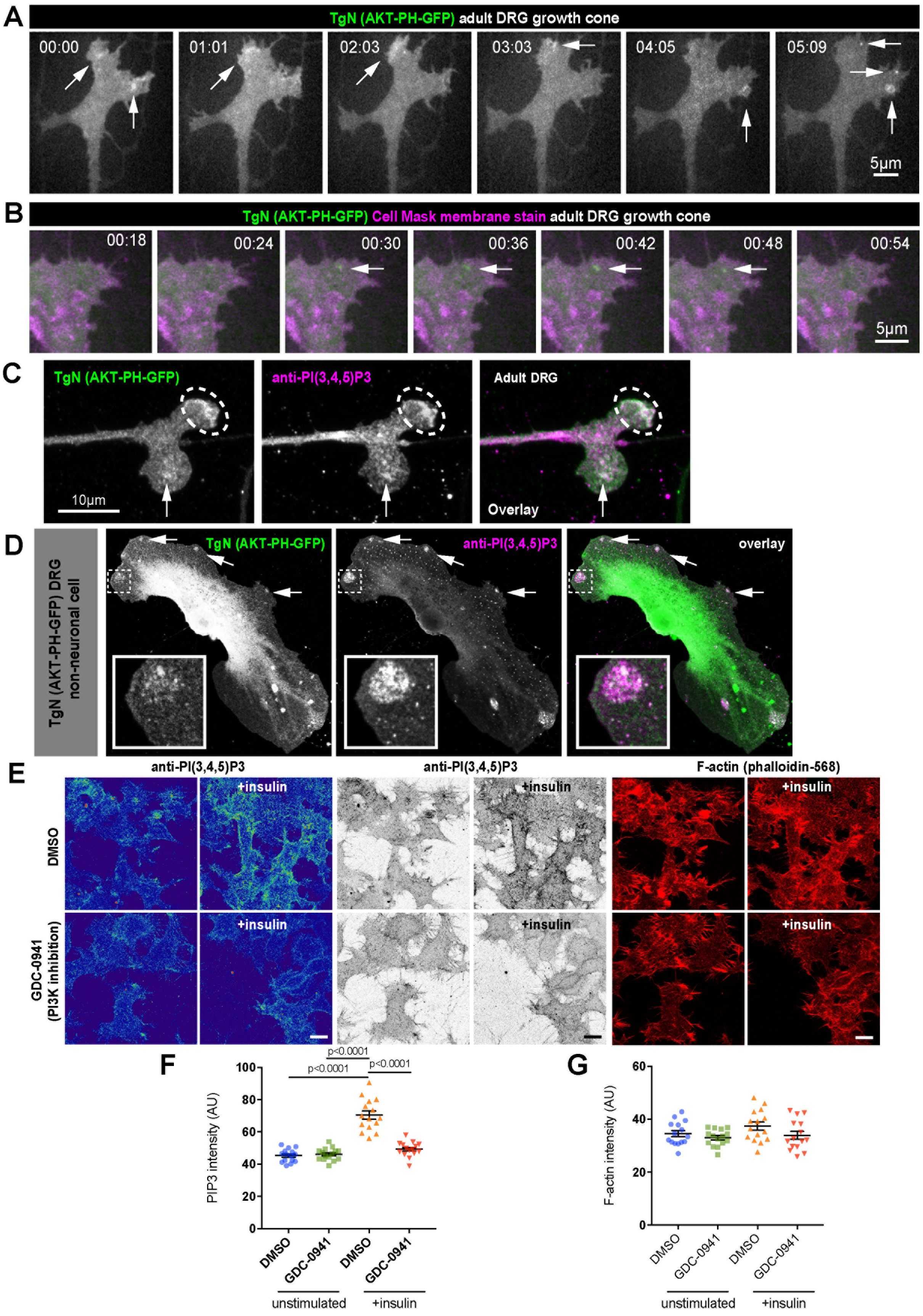
A
TgN (AKT-PH-GFP) adult DRG growth cone
00:00
01:01
02:03
03:03
04:05
05:09
5μm
B
TgN (AKT-PH-GFP) Cell Mask membrane stain adult DRG growth cone
00:18
00:24
00:30
00:36
00:42
00:48
00:54
5μm
C
TgN (AKT-PH-GFP)
anti-PI(3,4,5)P3
Adult DRG
10μm
Overlay
D
TgN (AKT-PH-GFP) DRG non-neuronal cell
TgN (AKT-PH-GFP)
anti-PI(3,4,5)P3
overlay
E
anti-PI(3,4,5)P3
anti-PI(3,4,5)P3
F-actin (phalloidin-568)
+insulin
DMSO
GDC-0941 (PI3K inhibition)
F
PIP3 intensity (AU)
p<0.0001
p<0.0001
p<0.0001
DMSO
GDC-0941
DMSO
GDC-0941
unstimulated
+insulin
G
F-actin intensity (AU)
DMSO
GDC-0941
DMSO
GDC-0941
unstimulated
+insulin

**Figure S2.**

**Figure S2. Validation of a protocol for detecting PI(3,4,5)$P_3$ in neuronal membranes.**

(A) Time-lapse images (single confocal section imaged by spinning disc microscopy) of a DRG growth cone cultured from adult AKT-PH-GFP mice. Arrows point to hotspots / regions of increased fluorescence indicative of AKT-PH recruitment. See also associated Movie 1.

(B) Time lapse Time-lapse images (single confocal section imaged by spinning disc microscopy) of a DRG growth cone cultured from adult AKT-PH-GFP mice, stained with cell-mask orange (magenta colour) to detect the membrane. Arrows indicate a dynamic region of AKT-PH-GFP recruitment that does not label for membrane aggregation. See also associated Movie 2.

(C) Adult DRG growth cone cultured from AKT-PH-GFP mice, fixed for PIP3 immobilisation (see methods section), and labelled with an antibody to PI(3,4,5)P3 (magenta). Arrows and dotted circles indicate colocalization.

(D) Non-neuronal cell from a dissociated DRG culture from AKT-PH-GFP mice, fixed for PIP3 immobilisation (see methods section), and labelled with an antibody to PI(3,4,5)$P_3$ (magenta). Arrows indicate colocalization. Inset highlights colocalization at a large region of AKT-PH-GFP recruitment.

(E) N1E neuroblastoma cells stimulated with insulin and labelled for PI(3,4,5)P3 in the presence or absence of the pan PI3K inhibitor GDC-0941. Cells were co-labelled for F-actin to show cell density.

(F) Quantification of PI(3,4,5)P in N1E neuroblastoma cells stimulated with insulin and in the presence or absence of the pan PI3K inhibitor GDC-0941. Insulin leads to an increase in PI(3,4,5,)P3 which is not detected in the presence of GDC-0941.

(G) Quantification of F-actin in N1E neuroblastoma cells stimulated with insulin and labelled for PI(3,4,5)$P_3$ in the presence or absence of the pan-PI3K inhibitor GDC-0941.

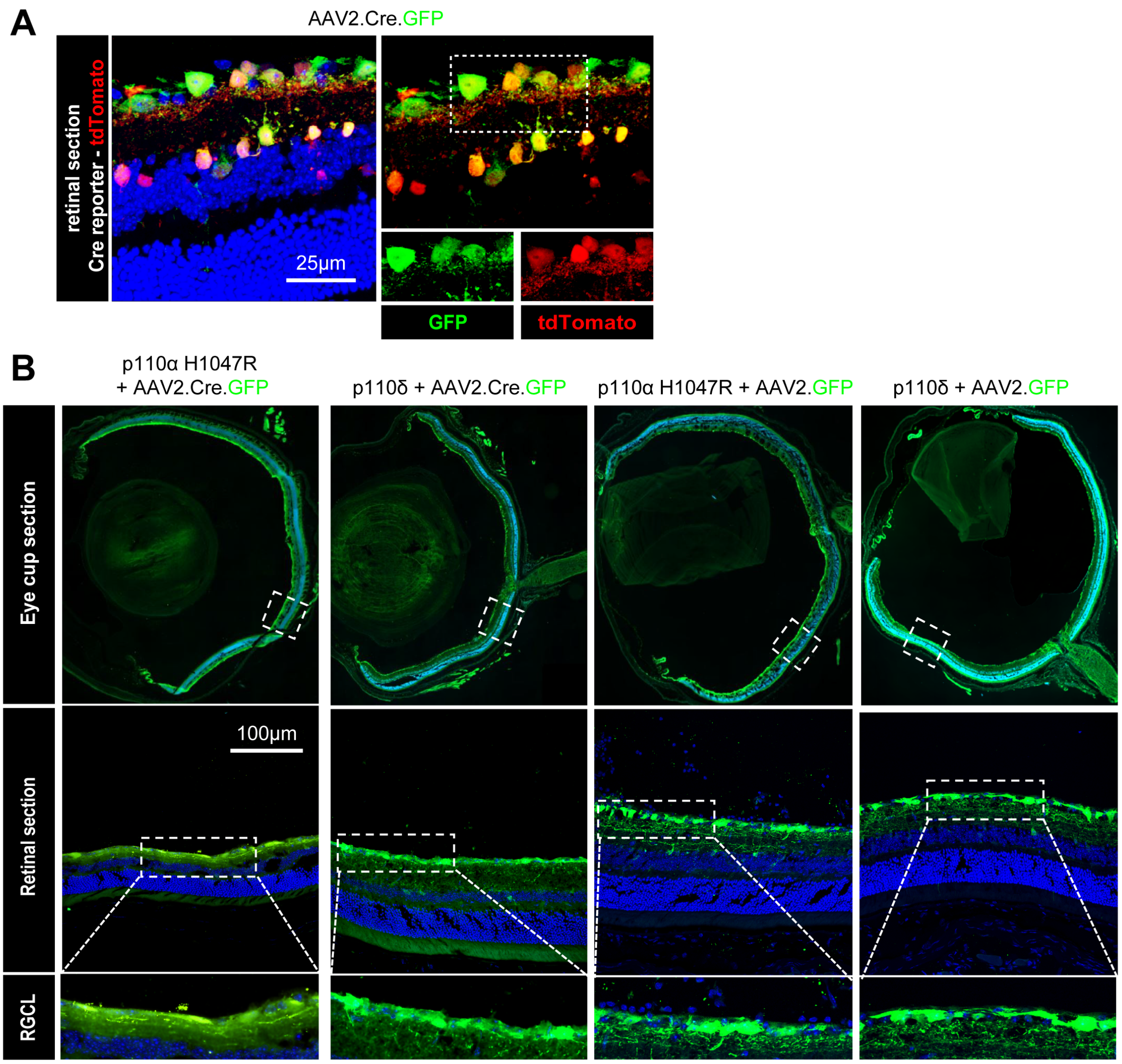


**Figure S3. Validation of recombinant adeno-associated viral vectors and targeting of RGC neurons.**

(A) Representative image showing expression of AAV2.Cre.GFP and co-localisation with TdTomato in Rosa26 Cre-reporter mice 2 weeks after injection of virus.

(B) Representative images showing expression of AAV2 vectors in eye cup- and retinal sections in the four treatment groups as indicated, 2 weeks after injection of virus. Images highlight expression in RGC layer (lower panels). Blue colour is DAPI nuclear staining.

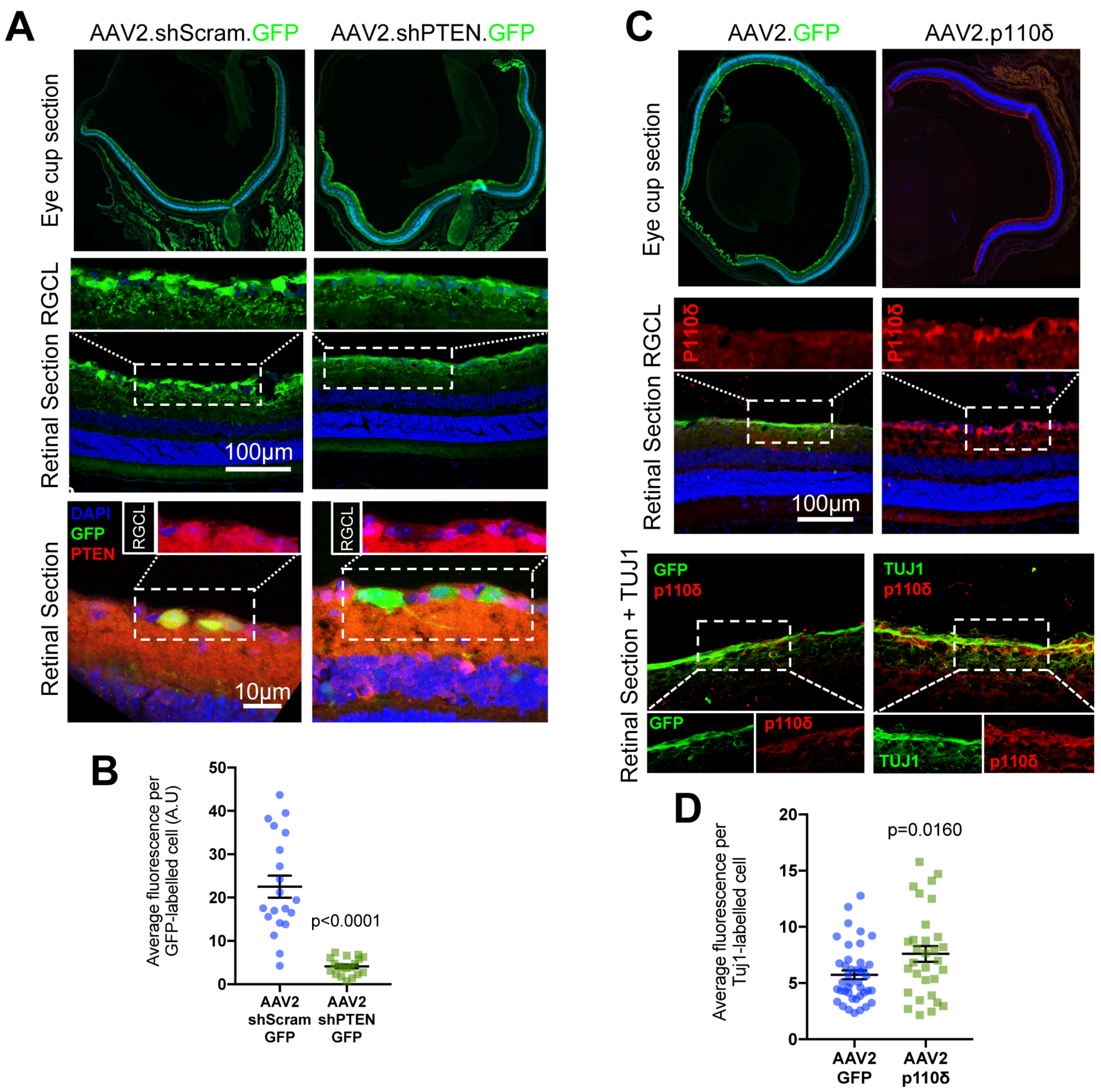


**Figure S4. Confirmation of AAV-mediated PTEN reduction and AAV transduction of p110δ in the retina**.

(A) Representative confocal images showing expression of AAV2.shScram.GFP (control) and AAV2.shPTEN.GFP in eye cup- and retinal cross-sections. Lower panels show a reduction in PTEN immunofluorescence in GFP positive cells in the RGC layer 2 weeks after injection with AAV2.shPTEN.GFP.

(B) Quantification of the PTEN immunofluorescence in transduced RGCs. Error bars are s.e.m. P values indicate statistical significance as measured by Students t-test.

(C) Representative images showing expression of AAV2.GFP and AAV2.p110δ in eye cup- and retinal sections 2 weeks after AAV injection. Retinae from eyes injected with AAV2.p110δ (right hand panels) were labelled for TUJ1 (green) and p110δ (red).

(D) Quantification of the p110δ immunofluorescence in RGCs transduced with AAV2.p110δ. Error bars are s.e.m. P values indicate statistical significance as measured by Students t-test.

### Movie Legends

**Movie 1.** DRG growth cone imaged by confocal spinning disc microscopy cultured from adult AKT-PH-GFP mice. Exhibiting hotspots / regions of increased fluorescence indicative of AKT-PH recruitment.

**Movie 2.** DRG growth cone from adult AKT-PH-GFP mice, stained with cell-mask orange to detect the membrane. Arrows indicate a dynamic region of AKT-PH-GFP recruitment that does not label for membrane aggregation (single confocal section imaged by spinning disc microscopy). Upper left panel is AKT-PH-GFP, upper right panel is cell-mask orange.

**Movie 3.** DIV 14 cortical neuron expressing p110δ and GFP regenerating after laser axotomy.

**Movie 4.** DIV 14 cortical neuron expressing GFP failing to regenerate after laser axotomy.

### Supporting Materials and Methods

#### Mouse Strains

The generation of the PIP3 reporter mouse GFP-AKT-PH has been previously described (Nishio et al, 2007) and was a gift from Dr Len Stephens (Babraham Institute, UK). Male and female mice were used dependent on litters available with equal distributions across experiments. Rosa26 p110α$^{H1047R}$, Rosa26 p110δ were generated according to a previously described protocol (Nyabi et al, 2009). Briefly, cDNA sequences for human p110α or p110δ were cloned into the pENTR D-TOPO vector (ThermoFisher) to allow cloning into Gateway-compatible vectors. The H1047R point mutation was introduced by site-directed mutagenesis into PIK3CA using the Quickchange II XL kit (Stratagene #200521) and the primers Quickchange H1047R forward 5'-catgaaacaaatgaatgatgcacgccatggtggctggacaac-3', Quickchange H1047R reverse 5'-gttgtccagccaccatggcgtgcatcattcatttgtttcatg. Presence of the mutation was confirmed by sequencing prior to cloning into the destination vector. PIK3CD (p110δ) was a gift from Roger Williams (MRC Laboratory for Molecular Biology, UK). PIK3CA (p110α) was a gift from Bert Vogelstein (Addgene plasmid #16643 ; http://n2t.net/addgene:16643; RRID:Addgene_16643) (Samuels et al, 2004). The Gateway®-compatible pROSA26 destination vector (pROSA26-DV1) was a gift from Jody Haigh

(University of Manitoba, Canada) (Nyabi et al, 2009). Targeting constructs harbouring sequences for $p110\alpha^{H1047R}$ or p110δ were used to electroporate Bruce4 C57BL/6 ES cells. Selection of targeted clones was undertaken with G418, making use of the neomycin resistance selection cassette. Positively targeted clones were identified by Southern blotting. Positively targeted clones were microinjected into C57BL/6J-Tyrc-2J blastocysts by Babraham Institute Gene Targeting Facility (Babraham Institute, UK). These blastocysts were transferred to the oviducts of time-mated pseudopregnant foster mothers. The progeny were assessed for their degree of chimerism by coat colour and chimeric males mated to white C57BL/6J-Tyrc-2J females. Black progeny were tested for their correct incorporation of the Human PI3K genes at the Rosa26 locus by PCR using the following primers:

| | | | |
|---|---|---|---|
| Rosa26 Hs p110 | ROSA26 F1 AMQ | GGCTCAGTTGGGCTGTTTTG | WT allele = 359 bp<br>KI allele = 603 bp |
| | ROSA26 R2 AMQ | TCTGTGGGAAGTCTTGTCCC | |
| | ROSA26 loxP AMQ | GTGGATGTGGAATGTGTGCG | |
| Hs p110α | Hs PIK3CA #5 intra fwd | TTGATCTTCGAATGTTACCT | KI allele = 813 bp |
| | S2rev | CTTGATATCGAATTCCGCCCC | |
| Hs p110δ | Hs PIK3CD #5 intra fwd | GTACTCCGTTCAGACACCAT | KI allele = 742 bp |
| | S2rev | CTTGATATCGAATTCCGCCCC | |
